# Innate Conformational Dynamics Drive Binding Specificity in Anti-Apoptotic Proteins Mcl-1 and Bcl-2

**DOI:** 10.1101/2022.06.10.495660

**Authors:** Esther Wolf, Cristina Lento, Jinyue Pu, Bryan C. Dickinson, Derek J. Wilson

## Abstract

The structurally conserved B-cell Lymphoma 2 (Bcl-2) family of proteins function to promote or inhibit apoptosis through an exceedingly complex web of specific, intrafamilial protein-protein interactions. The critical role of these proteins in lymphomas and other cancers has motivated a widespread interest in understanding the molecular mechanisms that drive specificity in Bcl-2 family interactions. However, the substantial structural similarity amongst Bcl-2 homologues has made it difficult to rationalize the highly specific (and often divergent) binding behavior exhibited by these proteins using conventional structural arguments. In this work, we use millisecond hydrogen deuterium exchange mass spectrometry to explore shifts in conformational dynamics associated with binding partner engagement in Bcl-2 family proteins Bcl-2 and Mcl-1. Using this approach, we reveal that, specifically for Mcl-1, binding specificity arises largely from protein-specific dynamic modes that are accessed in the unbound state. This work has implications for exploring the evolution of internally regulated biological systems composed of structurally similar proteins, and for the development of drugs targeting Bcl-2 family proteins for promotion of apoptosis in cancer.

**General Interest Statement:** This work reveals how a group of proteins, which are highly similar in structure, can form a complex web of highly specific protein-protein interactions that drive programmed cell death (apoptosis) in cancer.

## 1 INTRODUCTION

The mitochondrial outer membrane permeabilization (MOMP) pathway is an essential mechanism of programmed cell death (apoptosis) and is regulated through a complex web of interactions among Bcl-2 family proteins among other mechanisms. The Bcl-2 family encompasses many members that share Bcl-2 homology regions BH1-4: Pro-apoptotic multidomain proteins (Bak and Bax), pro-apoptotic BH3-only sensitizers (Bad, Noxa, etc.), pro-apoptotic BH3-only activators (tBid, Bim, etc.), and anti-apoptotic multidomain proteins (Bcl-2, Mcl-1, Bcl-X_L_, *etc*.).^1^

A brief snapshot of this intricate pathway begins when truncated Bid (tBid) interacts with Bak and Bax to facilitate their oligomerization within the mitochondrial membrane.^2,3^ This initiates an, irreversible commitment to apoptosis as cytochrome *c* effuses from the intermembrane space, apoptosomes (Apaf-l/cytochrome *c*/caspase-9) are formed, and subsequent caspase cascades lead to cellular destruction.^4^ However, anti-apoptotic proteins Bcl-2 and Mcl-1 can sequester tBid from this pathway and prevent it altogether. As such, the “ Dance Towards Death”, artfully named by Kale, Osterlund, and Andrews (2018)^1^, involves exchanging pro-and anti-apoptotic partners *via* heterodimerization at a conserved hydrophobic BH3-binding groove.

The BH3-binding groove exists on the surface of multidomain family members including Bak, Bax, Bcl-2, and Mcl-1. Notably, despite only sharing 33% sequence identity in their binding grooves,^5,6^ anti-apoptotic members Bcl-2 and Mcl-1 have nearly superimposable 3D structures: PyMOL backbone C_α_ alignment of Bcl-2 (PDB IG5M) and Mcl-1 (PDB 2MHS) for 106 core atoms yields an RMSD of 2.275 Å.^7–9^ Bcl-2 and Mcl-1 strongly interact with BH3-only proteins tBid and Bim; however, at physiological concentrations, only Mcl-1 can bind Noxa and only Bcl-can bind Bad. This poses an intriguing question about what molecular mechanisms can enable such unique binding selectivities in such structurally similar proteins.

Mass spectrometry provides a robust toolbox for elucidating molecular mechanisms including ‘Native’ ElectroSpray Ionization Mass Spectrometry (ESI-MS) to observe ligand binding, complexation and protein folding behaviour, Ion Mobility Separation Mass Spectrometry (IMS-MS) to analyze protein size/shape, and Hydrogen-Deuterium Exchange Mass Spectrometry (HDX-MS) to monitor structural dynamics and binding footprints with sub-molecular structural resolution. In particular, ‘time-resolved’ ElectroSpray Ionization MS (TRESI-HDX), which uses millisecond-to-low-second deuterium labeling times, can characterize subtle shifts in conformational dynamics that accompany protein complexation. These data can reveal binding sites, allostery, and provide structural/dynamic rationales for functional properties including binding specificity.

In this work, soluble Bcl-2 and Mcl-1 were examined using the mass spectrometry / hydrogen deuterium exchange toolbox to understand the molecular mechanisms that underly their distinct binding selectivities.^8,10^ Ion Mobility Spectrometry (IMS) showed that despite being a smaller protein construct by mass, Mcl-1 (18.05 kDa) exhibited a longer drift time than Bcl-2 (20.25 kDa), agreeing with previous classical structural studies indicating that Mcl-1 has a broader binding pocket.^9,11^ Binding studies using time-resolved HDX revealed that Bcl-2 and Mcl-1 undergo distinctive shifts in their conformational ensembles that are unique to each protein and occur regardless of whether they are interacting with a partner that binds one protein or both. From this we conclude that binding specificity in these Bcl-2 family proteins is driven not only by the charge compensation mechanisms proposed previously, but also by specific dynamic modes that are exhibited predominantly in the unbound state.

## 2 RESULTS

To express solution-stable recombinant Bcl-2 and Mcl-1 for *in vitro* study, optimized sequences representing the Bcl-2 homology core and purification strategies published by Petros et al. (2001) and Lee et al. (2016) were used.^8,10^ Petros et al. (2001) were the first to obtain a solution structure of Bcl-2. They based their solution-stable construct on Bcl-X_L_ which had been the first structure solved from the Bcl-2 family in 1997.^12^ By replacing the Bcl-2 ∼60 amino acid loop with the shorter loop of Bcl-X_L_, they were able to reduce disorder and lower the isoelectric point of the protein from near neutral pH to about pH 5.0.^8^ Previous publications indicated that Bcl-X_L_ retained function without it’s unstructured loop.^13^

Although attempts were made to work with purified wildtype Bcl-2 and Mcl-1, both displayed a propensity for aggregation and precipitation in MS-compatible solutions and subsequent clogging of sub-millimeter diameter capillaries used for delivery of sample to the mass spectrometer. This can be attributed to their transmembrane tails, large, disordered regions, and hydrophobic binding grooves. Consequently, all experiments discussed here use Mcl-1 (172-327) with an N-terminal SGS artifact after thrombin cleavage for GST tag removal and Bcl-2 (1-34, 35-50 Bcl-X_L_, 93-207) with a C-terminal His tag. The sequences used can be found in Table S1.

### 2.1 Native Mass Spectrometry

Previously, a review by Kale, Osterlund, and Andrews (2018) compiled over 30 publications which reported on the binding affinities of Bcl-2 family members using various biological assays including isothermal titration calorimetry (ITC), surface plasmon resonance (SPR), and fluorescence polarization (FP).^1^ Table 1 is adapted from their work and lists the Bcl-2 and Mcl-1 dissociation constants (K_D_) with BH3-only peptides Bim, Bid, Bad, and Noxa. From this work, Bcl-2 was expected to interact with Bid (50-10,000 nM K_D_), Bad (10-50 nM K_D_), and Bim (<10 nM K_D_), whereas Mcl-1 was expected to interact with Bid (<10 nM K_D_), Bim (<10 nM K_D_), and Noxa (10-100 nM K_D_). To confirm this and obtain the optimal molarity ratio of protein and BH3 peptide for the saturated “ bound” protein state, native electrospray ionization mass spectrometry (ESI-MS) was employed.

**Table 1.**
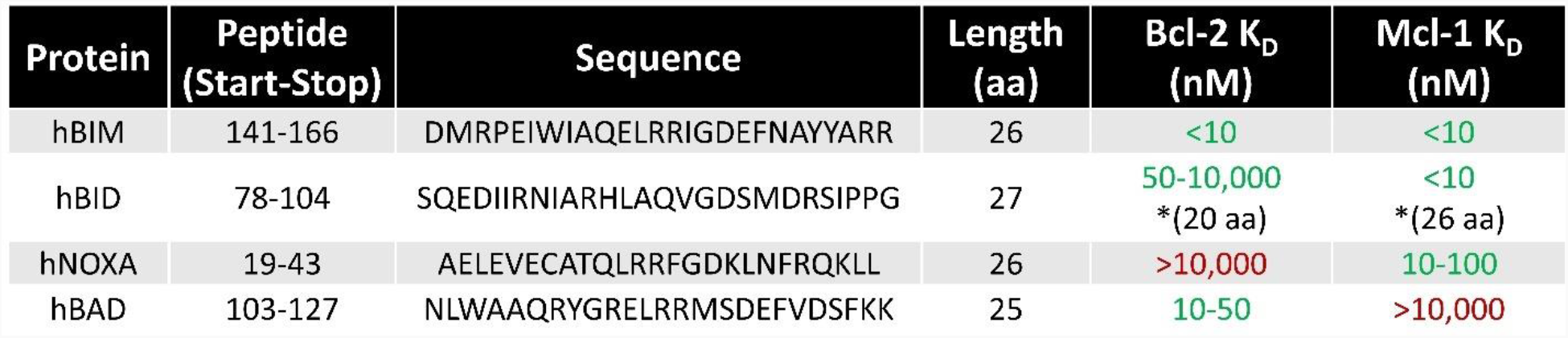
Dissociation Coefficients Reported for Select BH3 Peptides with Bcl-2 or Mcl-1

Native mass spectra are shown in Figure 1, starting with the unbound Bcl-2 and Mcl-1 spectra in panels A and B, respectively. Both proteins exhibited a similar, narrow charge distribution predominated by the 9+ and 8+ charged state, and minimally populated by 10+ and 7+. For Bcl-2, the four m/z charge peaks correspond to 2013.19, 2236.84, 2516.24, and 2875.60, whereas for Mcl-1 the four peaks correspond to m/z of 1806.41, 2006.93, 2257.64, and 2579.86. Using ESIprot deconvolution, the average masses of Bcl-2 and Mcl-1 was 20121.69 ± 0.85 Da and 18051.88 ± 1.04 Da, respectively.^14^ Although Mcl-1 matched its theoretical mass (18053.47 Da) based on its primary sequence, Bcl-2 (20253.53 Da) was off by 131.84 Da, which could be attributed to N-terminal methionine loss, a hypothesis supported by the N-terminal peptide later observed in HDX-MS experiments (shown in Figure 3, peptide 2-13).

**Figure 1.**
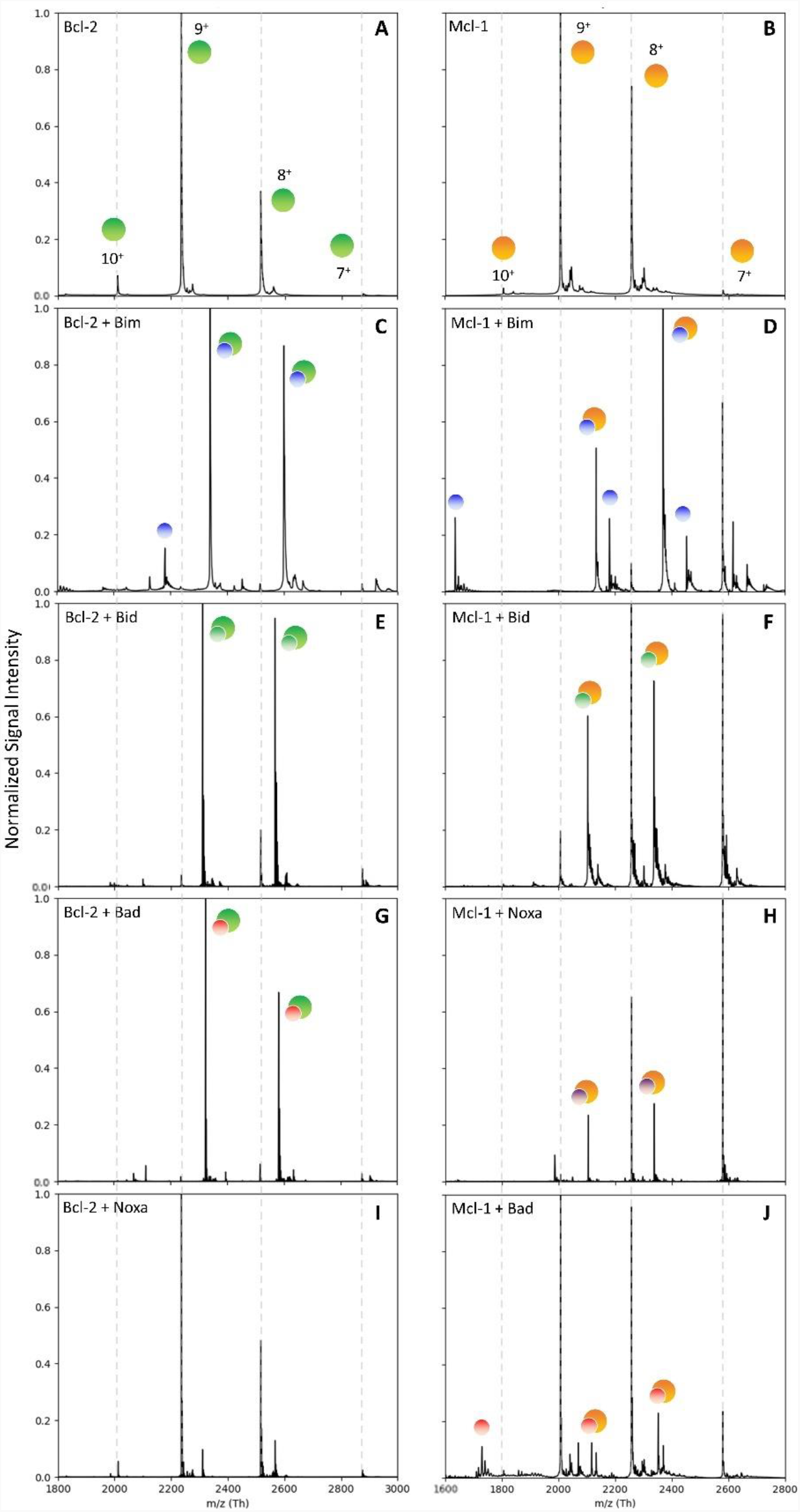
Native Mass Spectra of Bcl-2 and Mcl-1 bound to BH3 Peptides. A) Bcl-2; B) Mcl-1; C) Bcl-2 + Bim: 2340.27 (10+) and 2600.20 (9+); D) Mcl-1 + Bim: 2133.45 (10+) and 2370.37 (9+); E) Bcl-2 + Bid: 2310.87 (10+) and 2567.50 (9+); F) Mcl-1 + Bid: 2103.96 (10+) and 2337.68 (9+); G) Bcl-2 + Bad: 2323.60 (10+) and 2581.74 (9+); H) Mcl-1 + Noxa: 2104.39 (10+) and 2337.99 (9+). I) Bcl-2 + Noxa: no bound peaks; J) Mcl-1 + Bad: 2116.81 (10+) and 2351.92 (9+).

Next, 5 μM Bcl-2 or Mcl-1 was incubated with each BH3 peptide at a 1:1, 1:2, 1:4, and 1:6 molar ratio. As shown in figure 1C-H, a ratio of 1:6 generated saturated “ bound” native spectra for known interactors, and this was used as the optimal ratio for all subsequent HDX experiments (Figure 2). Dotted lines from figure 1A-B indicate the unbound peaks for Bcl-2 and Mcl-1. For Bcl-2 + Bim (23392.68 Da), the bound *m/z* peaks correspond to 2340.27 (10+) and 2600.20 (9+); for Bcl-2 + Bad (23226.26 Da), the bound *m/z* peaks correspond to 2323.60 (10+) and 2581.74 (9+); and for Bcl-2 + Bid (23098.53 Da), the bound *m/z* peaks correspond to 2310.87 (10+) and 2567.50 (9+). The experimental masses of the complexes fell within 2.6 Da of the expected masses of 23391.38 Da, 23225.20 Da, and 23096.01 Da, respectively. On the other hand, for the Mcl-1 + Bim complex (21324.35 Da), the *m/z* peaks correspond to 2133.45 (10+) and 2370.37 (9+); for Mcl-1 + Bid (21029.79 Da), the bound *m/z* peaks correspond to 2103.96 (10+) and 2337.68 (9+); and for Mcl-1 + Noxa (21033.33 Da), the bound peaks correspond to 2104.39 (10+) and 2337.99 (9+). Here, the maximum mass deviation was about 4 Da from the expected masses of 21324.35 Da, 21026.20 Da, and 21030.34 Da, respectively. Interestingly, in the Bcl-2 ‘bound’ spectra, peaks corresponding to the complex dominate, whereas in the Mcl-1 spectra, binding appears to induce a shift to lower charge, even in peaks for *m/z* peaks corresponding to the unbound protein. This may arise from loss of the Mcl-1/BH1 peptide complex in the gas phase accompanied by charge stripping by the departing peptide, resulting in a significant fraction of the lower-charge ‘unbound’ peak intensity being attributable to protein that was originally ‘bound’ in solution.

**Figure 2.**
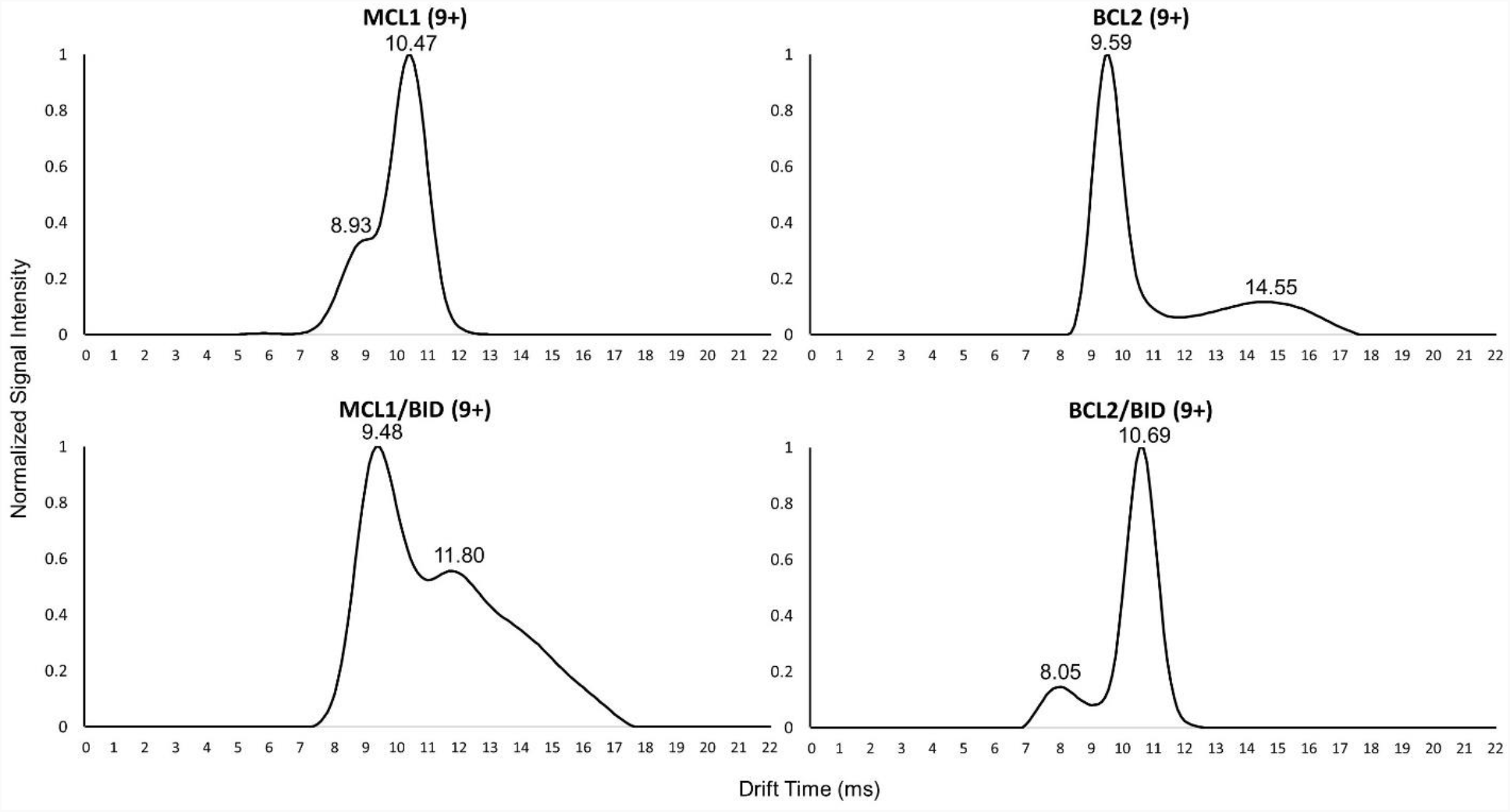
Ion Mobility Chromatograms of Bcl-2 and Mcl-1. The relative signal intensity as a function of drift time (milliseconds) for Mcl-1 (top left), Bcl-2 (top right), Mcl-1 + Bid (bottom left), and Bcl-2 + Bid (bottom right) is shown for their 9+ charge state.

No interaction was detected for Bcl-2 + Noxa (Figure 1I), which is consistent with what is known about Bcl-2 specificity (Noxa is an Mcl-1 specific binder). Minor peaks corresponding to Mcl-1 + Bad (a Bcl-2 specific binder) were observed (Figure 1J), suggesting the possibility of weak binding. However, Bad did not induce the charge reduction effect observed for all other binders of Mcl-1, indicating that the maximum ‘bound’ fraction can be estimated, based on the intensities of the ‘bound’ peaks relative to the unbound peaks, to be no more than 25%. Note that this estimate excludes the very real possibility that some or all of the observed ‘bound’ peaks arise from non-specific complexation or adduction, which is a common phenomenon in ESI mass spectrometry.^15^

### 2.2 Ion Mobility Mass Spectrometry

Ion Mobility Separation Mass Spectrometry (IMS-MS) was used to examine the gas phase conformations that populated the native ESI-MS spectra of Bcl-2 and Mcl-1. In Figure 2, the normalized intensity of signal (ion count) is recorded as a function of the ion mobility drift time in milliseconds, with lower drift times corresponding generally to smaller globular size (*via* the collisional cross section) for a given charge state. The 9+ charge state was selected because it is well populated in both the unbound and bound state spectra for both proteins (and comparing structures of the same charge reduces the complexity of interpreting IMS data).

The dominant drift time peak for unbound Mcl-1 was centered on 10.47 ms whereas the dominant peak for Bcl-2 was 9.59 ms (Figure 2, top row). This is the opposite of what might be expected intuitively, since the Bcl-2 construct used in this study is 15 residues longer than the Mcl-1 construct. However, the impact of this additional sequence on the collisional cross section (which is ultimately what IMS measures) could easily be subsumed by differences in how the protein is packed overall. In any event, based on the dominant peaks, it appears that Mcl-1 has a somewhat larger cross section than Bcl-2. Both proteins also exhibit minor gas phase configurations in the unbound state, corresponding a ‘compact’ structure in the case of Mcl-1 (shoulder at 8.93 ms) and an ‘extended/disordered’ structure for Bcl-2 (broad low peak at 14.55 ms). Additional IMS plots for the 10+ charge state can be found in Figure S1.

Upon complexation with the Bid BH3 peptide, Mcl-1 (2104 *m/z*) and Bcl-2 (2311 *m/z*) undergo distinct changes in their gas phase configurations. For relatively small proteins like Mcl-1 and Bcl-2, complexation with a large peptide such as Bid BH3 (2976.32 Da) could cause an increase or decrease in the CCS, depending on how the incoming peptide packs onto the structure, and the extent to which the conformation ‘tightens’ as new bonds are formed with the binder. In this case, complexation appears to have had the opposite effects on Mcl-1 and Bcl-2 drift times, resulting in a ‘tightening’ of the structure for Mcl-1 (10.47 ms – 9.48 ms), and a larger cross section for Bcl-2 (9.59 ms – 10.59 ms). At the same time, the minor peaks have switched places relative to the main peaks, so that bound Mcl-1 is exhibiting a new extended configuration (11.80 ms) and Bcl-2 is exhibiting a new compact state (8.05 ms) upon binding. Taken together, these IMS data suggest significantly different binding modes for Mcl-1 and Bcl-2 in their interaction with Bid, however, any characterization of these differences from IMS alone would be, at best, highly speculative, and there is no guarantee that our gas phase observations directly reflect the process in solution. Our next step was therefore to undertake a TRESI-HDX analysis.

### 2.3 Hydrogen-Deuterium Exchange Mass Spectrometry

Time-resolved HDX was performed using an adjustable mixer composed of two concentric sub-millimeter diameter capillaries coupled to a pepsin-functionalized microfluidic chip. This method for ‘bottom-up millisecond hydrogen deuterium exchange’ has been described previously.^16–18^ It enables ‘segment-averaged’ (peptide-level) measurements of deuterium uptake with millisecond-second labeling times, which can probe subtle shifts in conformational dynamics resulting from complexation. Here, the HDX data are presented differentially, meaning that deuterium uptake in unbound protein is subtracted from uptake in the peptide-bound protein (Figure 3). As a result, bars with negative values indicate that deuterium uptake has decreased in the corresponding region as a result of complexation. HDX difference profiles were acquired for Bcl-2 and Mcl-1 upon complexation with peptides corresponding to the BH3 peptides of Bim, Bid, Bad and Noxa. For normalized relative uptake (%) of each state, please see supplementary figures S2-S9.

**Figure 3.**
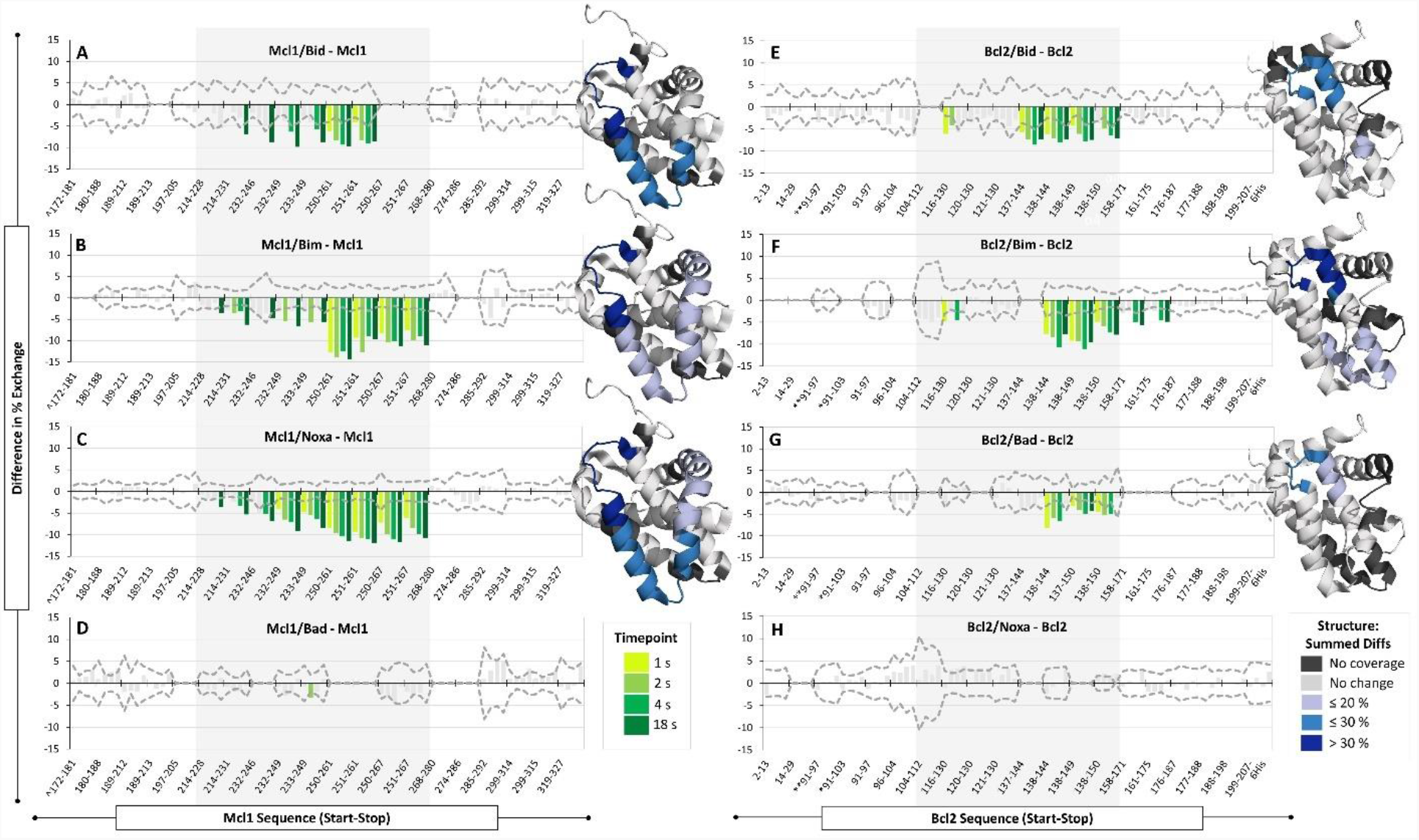
Difference in % Exchange of Complex versus unbound Bcl-2 and Mcl-1. Time-resolved HDX-MS was conducted at 1 s, 2 s, 4 s, and 18 s labeling times for Mcl-1 or Bcl-2 with and without incubation of Noxa, Bid, Bim, or Bad BH3 peptides. Bar plots represent a differential analysis of the HDX data, corresponding to the ‘unbound’ exchange profile subtracted from the ‘bound’ profile. Coloured bars indicate a statistically significant decrease in uptake with different shades of green corresponding to the 4 different labeling times. To be considered statistically significant, the ‘difference in % exchange’ had to exceed the propagated error (2σ, dashed line) by 1% on the ‘difference in % exchange’ scale. The peptides within the BH3 binding groove are highlighted in grey shading. The structures on the right are colour coded based on summed differences in % exchange and correspond to the following: no sequence coverage (black), no change (light grey), ≤ 20 % (light blue), ≤ 30 % (sky blue), > 30% (royal blue).

One of the disadvantages of online time-resolved HDX is that without liquid chromatographic separation, it often provides lower sequence coverage than the corresponding conventional timescale experiment. Sequence coverage varied considerably depending on the system in question, with the highest coverage corresponding to Bcl-2 + Bim (90%), the lowest to Bcl-2 + Noxa (62%), and an average of 81% across the entire dataset (sequence coverage and redundancy data are provided in Table S2 and the peptide list for each protein is provided in Tables S3 and S4). In every case, peptides located within the binding grooves of Bcl-2 and Mcl-2 were observed. The average peptide length was 10 amino acids, with segments ranging from 7 – 20 residues. Peptides in Bcl-2 involving chimeric sequence are indicated using an asterisk (*) to denote 39-50 of Bcl-X_L_ or a double asterisk (**) to denote 36-50 of Bcl-X_L_. In Mcl-1, the N-terminus peptide contains a thrombin cleavage site artifact (S) denoted by a circumflex (^).

Time-resolved HDX was performed at four timepoints: 1 s, 2 s, 4 s, and 18 s, which are represented in the Figure 3 differential bar plots as increasingly dark shades of green. The x-axes for these plots include all peptides detected for the corresponding protein. Thus, ‘blank’ segments in the bar plots indicate regions where peptides were detected in other bound states, but not the one in question. The dashed line on the bar graphs represents the propagated error and uses the two-fold standard deviation (2σ) of at least two replicates of three technical triplicates per state (*n* = 6). Bars that did not exceed this magnitude by more than 1% on the ‘difference in % exchange’ scale are coloured light grey to indicate that they were not considered statistically significant.

Figure 3 also highlights regions with significant changes in summed differences of % exchange, mapped onto NMR-derived structures of Mcl-1 (PDB 2MHS) and Bcl-2 (PDB 1G5M).^8,9^ Light-, sky-, and royal blue were used to illustrate the summed differences totalling ≤ 20 %, ≤ 30 %, and > 30%, respectively. Regions for which no peptides were obtained are denoted in black and regions for which there was no significant change are represented in grey.

For Mcl-1/Noxa (Figure 3C), persistent, large magnitude decreases were observed in peptides spanning 232-267 and late-appearing (but persistent) decreases were observed from 214-246. A nearly identical profile was observed for Mcl-1/Bim in Figure 3B. The Mcl-1/Bid profile (Figure 3A) also showed persistent decreases at 250-261 and late decreases at 232-249; however, some peptides corresponding to the 250-267 region could not be analyzed due to extensive signal overlap in the raw uptake data. For all three interactions with Mcl-1 (*i*.*e*., Noxa, Bid, and Bim), all significant changes occurred across the conserved binding groove at helices α3-α5. In the case of Mcl-1/Bad (Figure 3D), essentially no significant changes in deuterium uptake were observed across the entire protein, consistent with the expected lack of interaction.

For Bcl-2, the interaction with Bid resulted in persistent changes in a much narrower region corresponding to residues 137-150 (Figure 3E), which maps to an area within the Bcl-2 binding groove. A weak ‘signal’ is observed in the 116-130 region but is not observed in broadly overlapping peptides 120-130 and 121-130. The summed difference signal is therefore only plotted from 116-119 in the corresponding NMR structure. Similarly, for Bcl-2/Bim (Figure 3F), persistent decreases were observed at 138-150 together with a weak, inconsistent signal at 116-130, and a slowly developing decrease was observed in the 158-175 region. The Bcl-2/Bad complex in Figure 3G exhibited the weakest decreases in uptake, but these were all persistent and occurred in the 138-150 region in agreement with the other Bcl-2 complexes. Notably, in all 3 binding scenarios, the persistent decreases occurred at a localized region within the binding groove at the junction of helices α4-α5, corresponding to the BH1 domain that facilitates complexation in Bcl-2 and Mcl-1 *via* a conserved intermolecular salt-bridge. Bcl-2 did not exhibit any significant changes in deuterium uptake in the presence of Noxa (Figure 3H), which is consistent with the expected lack of interaction.

## 3 DISCUSSION

### 3.1 Native MS and IM-MS

The Bcl-2 and Mcl-1 constructs used here both consist of 7 α-helices: the predominantly hydrophobic helix α5 makes up the core of both proteins, which is surrounded by amphipathic helices α1-4 and α6-7.^19^ Distinctly, the α3 in Bcl-2 is considered a 3^10^-helix. Native MS revealed that despite their different masses, both recombinant Bcl-2 and Mcl-1 exhibit highly similar, narrow, low charge magnitude charge-state envelopes. This suggests that these proteins were ionized with the broadly comparable structural topology that is conserved within the anti-apoptotic Bcl-2 family members and is consistent with their known structures (Figure 4).^8,9,11^

**Figure 4.**
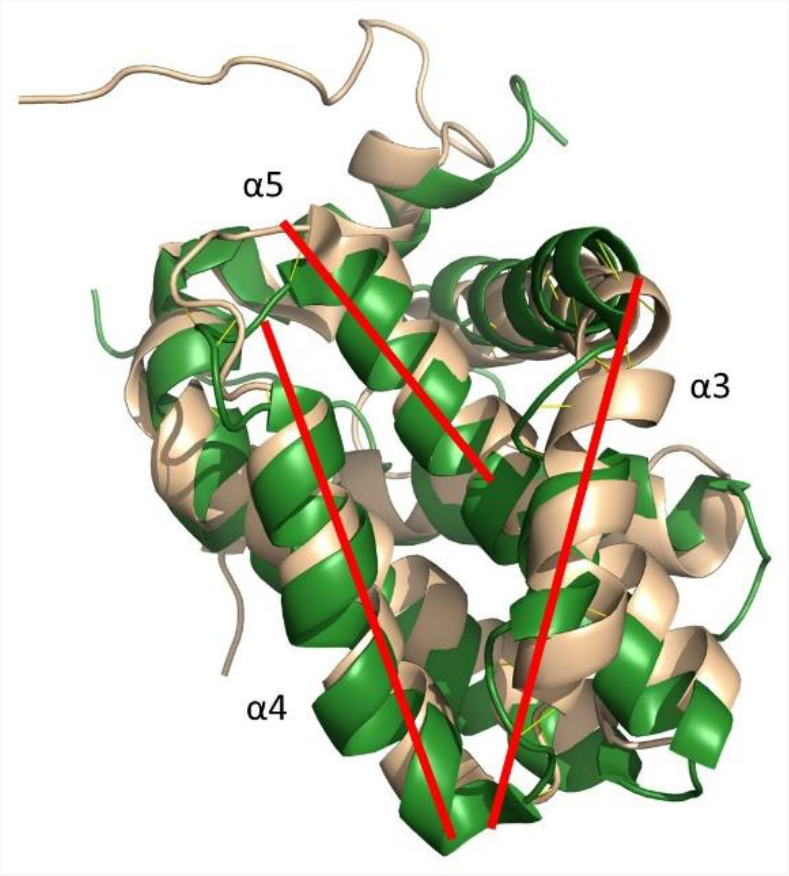
Backbone Aligned Mcl-1 and Bcl-2 Structure. Mcl-1 (yellow) and Bcl-2 (green) share the Bcl-2 homology core which includes the conserved BH3 binding groove composed of helices α3-α5 (red lines). The solution structures used here were 2MHS (Mcl-1) and 1G5M (Bcl-2).^8,9^

**Figure 5.**
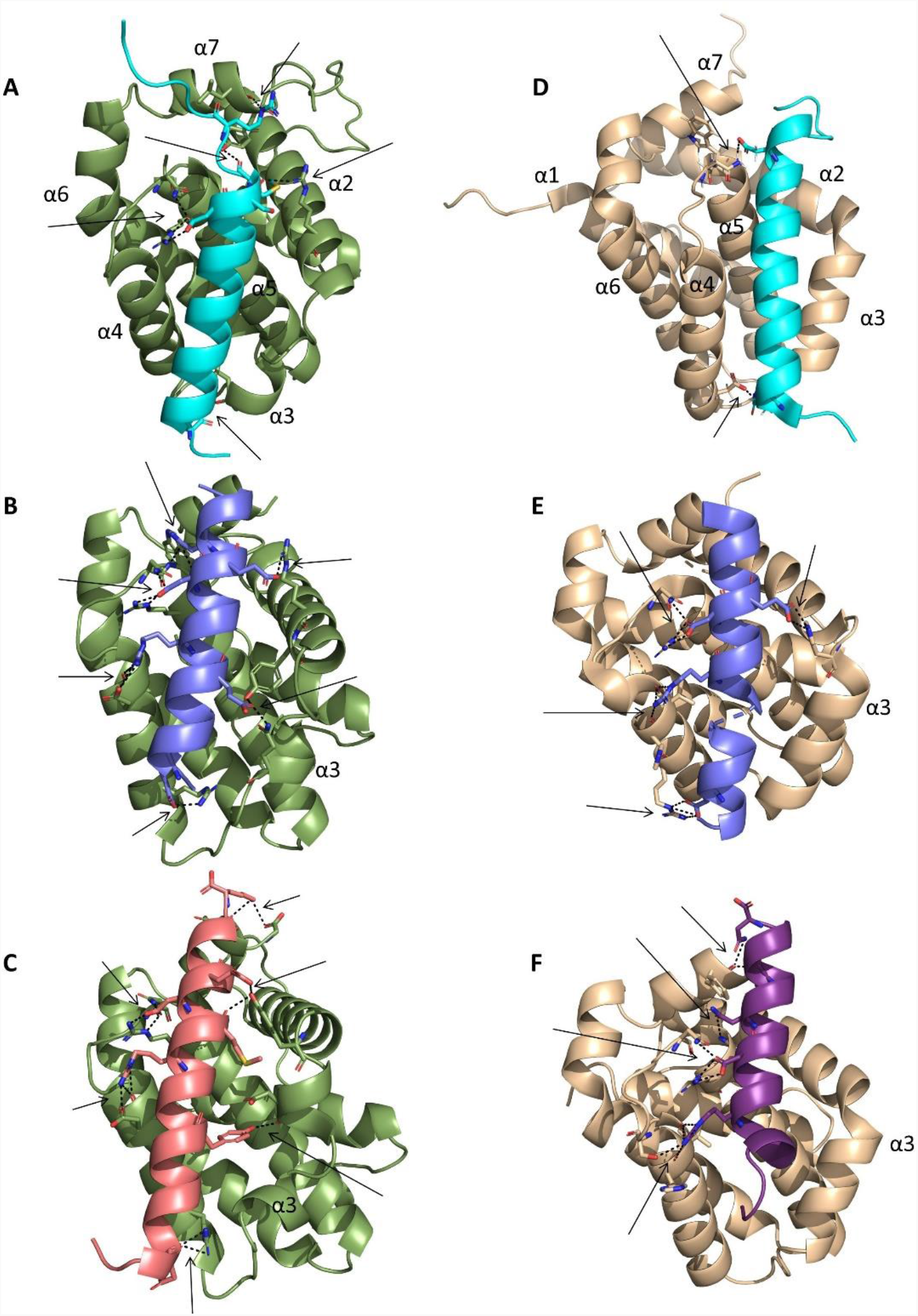
Homology Models and PDB Structures Displaying Intermolecular Bonding. Intermolecular bonds denoted in black dashed lines and emphasized with black arrows. Bcl-2 is green, Mcl-1 is tan-coloured. For the BH3 peptides, Bid is cyan, Bim is periwinkle, Bad is salmon, Noxa is violet.

However, the IMS data show a distinct difference in drift time, with Mcl-1 exhibiting longer drift times than Bcl-2. This observation is in agreement with previous work suggesting that Mcl-1 has a broader binding groove than other antiapoptotic members of the Bcl-2 family.^9^ The multimodal distributions in the IMS data also point to coexisting conformations in the gas phase that may reflect low-abundance conformational configurations in solution. In the case of Bcl-2, the additional conformation appears as a ‘larger’ (less folded) configuration that is structurally heterogeneous. The IMS data for Mcl-1 indicate that the dominant mobility peak is more extended than the minor population that appears as a shoulder, but retains a relatively homogeneous (i.e. ordered) conformational ensemble.

When complexed with Bid BH3, the dominant configuration of Mcl-1 exhibits a decreased drift time, which would be consistent with substantial penetration of the BH3 peptide into the binding groove coupled with an overall ‘tightening’ of the protein structure (Figure 2, left column). However, the dominant IMS peak is accompanied by a broad, substantially higher drift time peak which indicates that a significant fraction of the Mcl-1 population has undergone a conformational shift making the protein structure more extended and heterogeneous (or more susceptible to unfolding in the gas phase) upon complexation. While the dominant Mcl-1 + Bid IMS peak is relatively easy to rationalize in the context of complexation, it is not clear what binding effects might cause an increase in drift time, although in principle, both conformational disruption and binding without penetration into the binding grove are possibilities.

Conversely, Bcl-2 bound to Bid BH3 has a narrowly distributed, longer drift time peak that is indicative of a larger collisional cross section compared to unbound Bcl-2 (Figure 2, right column). This could imply that Bid binding does not involve deep penetration of the BH3 peptide into the Bcl-2 binding groove and does not involve substantial rearrangement of the protein structure, which would be consistent with most observations from conventional structural studies.^20^ However, it is as always rather difficult to draw unambiguous conclusions about processes occurring in solution from IMS data alone, and the appearance of a highly compact minor peak upon binding (Figure 3, bottom right) does not fit with this narrative. Some of these ambiguities can be addressed using time-resolved hydrogen deuterium exchange, which is carried out in solution and provides a higher degree of structural resolution.

### 3.2 Structure and Dynamics of Bcl-2 and Mcl-1

The basis of heterodimerization between pro-and anti-apoptotic family members has long been thought to be driven primarily by hydrophobic interactions.^1^ The BH3 groove in anti-apoptotic members is composed of α3-α5 which has four hydrophobic “ pockets”, denoted as P1-P4, and a conserved Arg residue at in the BH1 domain (α4-α5 loop).^21^ The BH3-only proapoptotic members have a conserved Asp (i+9) and four hydrophobic residues: H1 (i), H2 (i+4), H3 (i+7), and H4 (i+11) which line up to form the hydrophobic face of the amphipathic BH3 helix. The conserved Asp and Arg form an intermolecular salt bridge at BH1 domain of Bcl-2 and Mcl-1 (α4-α5 hinge DGV(T_Mcl1_)NWGR).^20^ However, these conserved interactions fail to explain why the anti-apoptotic proteins have distinct binding selectivities despite their highly similar function and topology.

Differential HDX revealed that Bcl-2 and Mcl-1 each exhibit a unique structural and dynamic behaviour during binding, regardless of the BH3 peptide that is bound. Mcl-1 undergoes a large, broadly distributed conformational change within the binding groove at α3-α5. To the best of our knowledge, Bcl-2 has never been studied by HDX-MS; however, in agreement with our findings, previously published work by Lee *et al*. (2016) reported broadly distributed decreases in deuterium uptake for Mcl-1+Bid at a 10 s HDX timepoint. Using LC-HDX-MS they detected high magnitude decreases at α3, α4, the C-terminal of α2, and N-terminal of α5, which is a more geographically constrained effect than we observe here (likely a result of the improved sensitivity of ms HDX measurements to subtle changes in dynamics).^10^ In contrast to Mcl-1, Bcl-2 exhibits a decrease in uptake that is localized specifically to the α4-α5 hinge (BH1 domain) with a small decrease at the C terminal end of α3 for two out of three binders.

Taken together, the HDX data indicate that Mcl-1 must undergo a substantial change in structure and/or dynamics to accommodate heterodimerization, whereas in Bcl-2, binding is driven less by dynamic shifts and more by specific interactions in the conserved BH1 region. One way of interpreting these results is that higher flexibility in the Mcl-1 binding pocket results in ‘specificity’ simply because it allows for the accommodation of larger BH3 ligands. However, the BH3 ligands used in this study are all similar in size. An alternative explanation arises from a recent study on the N-terminal domain of p53 whose binding specificity is modulated by phosphorylation at specific sites.^22^ In that case, the authors were able to demonstrate that conformational instability in the unbound state can be an enthalpic driver of complexation, and a mechanism for divergent binding specificity between protein states (in that case, unphosphorylated *vs*. phosphorylated p53 N-terminus). In this case, it may be that conformational instability in the Mcl-1 unbound state makes complexation thermodynamically favorable even in cases where other driving factors (*e*.*g*. hydrogen bonding, charge compensation *etc*.) are weaker. This may explain the apparent weak affinity of Mcl-1 for ‘non-binder’ Bad observed in the Native MS spectra.

To further explore the chemical interactions between the human BH3 peptides and Bcl-2 or Mcl-1, we used SWISS-MODEL^23^, an open-access protein homology-modeling software package, to visualize 3D structures of Bcl-2 or Mcl-1 with ‘bound’ BH3 peptides for which there are currently no co-crystal or NMR structures. In Bcl-2 homology modeling, Bcl-X_L_ was used as the template whereas the Mcl-1 Noxa model was based on a template of Mcl-1 bound to mouse NoxaB.

Overall, Bcl-2 was observed to form a higher number of peptide-protein interactions (i.e., salt-bridges and hydrogen bonds) compared to Mcl-1 in its interactions with all BH3 peptides. However, this general observation provides an example of the limits of a purely static structural analysis of these protein interactions because it offers no basis for specificity differences between Bcl-2 and Mcl-1 and also (incorrectly) suggests that Bcl-2 should bind all BH3 targets tested more tightly than Mcl-1. Nonetheless, the homology modelling approach was able to capture important interactions that were reflected in the HDX data. Specifically, the conserved salt bridge between anti-apoptotic BH3 Arg and pro-apoptotic Asp of the BH1 domain of the α4-α5 loop was detected in all interactions. In the case of Bcl-2 + Bid, additional interactions were observed at the C-termini of α2, α3, and α7. For Bcl-2 + Bim, interactions were observed at α2, α3, and α4, all of which were reflected in the HDX data. For Bcl-2 + Bad, there appears to be a restructuring at the α2-α3 region to enable two binding interactions in addition to salt bridges at the C-and N-termini of α4 and the C-terminus of α7. These α4 and α7 interactions were not detected by HDX. It is not clear if this is a result of a lack of sensitivity of HDX to these particular changes or an incorrect prediction from the homology modeling approach we are using here.

One unique difference for Mcl-1 interactions predicted by homology modelling was that none of the heterodimerizations involved α3. For Mcl-1 + Bim, interactions were formed at the α2-α3 loop, as well as the middle and C-terminal region of α4 (PDB 2PQK). For Mcl-1 + Bid, the only interaction outside of the conserved Arg-Asp salt bridge was another salt-bridge at the N-terminus of α4 (PDB 2KBW). As for Mcl-1 + Noxa, homology modeling revealed interactions at the C-termini of α4 and α7. An aspect of the experimental data that is supported by the homology models is the observation of an IMS drift time increase for Bcl-2 and a drift-time decrease for Mcl-1 upon complexation. This is reflected by the fact that in the Bcl-2 models, the BH3 peptide lies ‘flat’ across the binding grove, with little penetration into it. Conversely, in Mcl-1 models, the BH3 peptide is partially enclosed in the binding group by α3 and α4.

Taken together, our data point to a striking difference in how Mcl-1 and Bcl-2 interact with their targets, despite their high degree of structural similarity. Specifically, Bcl-2 interactions appear to be driven largely by charge compensation between the protein and its targets, with relatively subtle and localized changes in conformational dynamics upon binding. In contrast, Mcl-1 interactions are driven more by the transition from an unbound structure with fewer intramolecular hydrogen bonds to a bound structure with more intramolecular hydrogen bonds, and less by specific chemical interactions between the protein and its targets. As a result, Bcl-2 complexation is highly specific for BH3 segments that have charged residues in positions that are complementary to its binding grove while Mcl-1 complexation is specific for BH3 segments that can induce the needed rearrangement of Mcl-1 structure by insertion into the binding groove.

## 4 CONCLUSION

This study explores the basis for target specificity in the Bcl-2 protein family by focusing on two members, Bcl-2 and Mcl-1, that are structurally similar, but are known to exhibit different binding specificities. More fundamentally, though, we aim to shed light on the question of how structurally similar proteins can exhibit divergent specificities, a question that is relevant to a host of protein families composed of structurally similar homologues. In this case, our observation that Bcl-2 binding specificity appeared to be closely associated with the positioning of charged residues in the target is the one that arises naturally from conventional structural analyses and indeed, this has been extensively explored for Bcl-2 homologues.^24^ However, this explanation is insufficient to account for the intricacies of Bcl-2 family binding specificities, particularly as in the case of Bcl-2 and Mcl-1, where specificity profiles are partially overlapping. Such specificity profiles can only be fully understood by also accounting for other drivers of complexation, especially (in this case) conformational transitions that involve the formation of new intramolecular hydrogen bonds.

From the perspective of molecular evolution, these results provide a potential hypothesis as to how the specificity of interactions between proteins can be altered through a difficult-to-predict suite of mutations. In recent work, we demonstrated that the evolution of Mcl-1 and Bcl-2 specificity was ‘path-dependent’, implying that divergent evolutionary pathways generate unique and functionally inequivalent solutions (*i*.*e*., sets of mutations) to subsequent evolutionary challenges.^25^ The results of the current work reinforce the view that specificity is not dictated by specific amino acid substitutions at the interface but can also be driven by widely dispersed mutations that impact protein-wide conformational dynamics. Future efforts examining how alterations in protein dynamics impacted the evolution of functions within Mcl-1 and Bcl-2, and specifically whether there is an identifiable ‘branch-point’ ancestor whose evolution diverged into ‘charge-dominated’ and ‘dynamics dominated’ branches, could shed light on how natural protein functions emerged, and could also provide new approaches for engineering novel binding specificity in biomolecules.

From the perspective of practical outcomes from this type of analysis, a more complete understanding of binding specificity among structurally similar proteins is a foundation for the development of targeted therapeutics in many ‘challenging’ protein families, including the Bcl-2, GST, LXR and GPCR families among many others. Our hope is that further explorations of the dynamic drivers of binding specificity will encourage drug development targeting not just structures, but function-critical structural transitions.

## 5 MATERIALS AND METHODS

### 5.1 Materials

#### Protein Purification

Plasmids encoding Bcl-2 and Mcl-1 were purchased from Biobasic Inc. (Toronto, Canada) and constructed based on solution stable sequences optimized by the labs of SW Fesik and LD Walensky, respectively.^8,10^ Briefly, Bcl-2 was expressed in BL21(DE3) at 16°C for 18 hrs using pET28a (1-34 Bcl-2, 35-50 Bcl-XL, 93-207 Bcl-2) with a C-terminal 6His tag and purified using Ni^2+^ IMAC (GE Healthcare, Fastflow). Similarly, Mcl-1 was expressed in BL21(DE3) at 16°C for 18 hrs using pGEX-4T.1 encoding Mcl-1 (172-327) with an N-terminal GST-thrombin tag. GST tagged Mcl-1 was purified using GST-affinity chromatography (Glutathione-Sepharose resin by GE Healthcare, Fastflow). Subsequent GST-tag cleavage was carried out under rotation overnight at 4°C (Thrombin from bovine plasma, Sigma-Aldrich) followed by secondary GST-purification to remove free GST tag (untagged Mcl-1 collected in flowthrough). Both Bcl-2-6His and untagged Mcl-1 were gently concentrated/buffer exchanged into 50 mM Tris, 150 mM NaCl, 1 mM EDTA, pH 7.0 by spin-sized centrifugal filtration (Amicon, 10 kDa MWCO) at 1200 g, 4°C, 12 min cycles, resuspending by pipetting up-down between each cycle. For long-term storage at -80°C, glycerol was added to a final concentration of 25% and the sample was aliquoted such that no tube was ever thawed more than once. Protein identity was verified by 15% SDS-PAGE and intact-MS.

#### BH3-only Peptides

The following BH3-only peptides were synthesized and purchased from BioBasic Inc (Toronto, Canada): hBID (78-104) SQEDIIRNIARHLAQVGDSMDRSIPPG, hNOXA (19-43) AELEVE-CATQLRRFGDKLNFRQKLL, hBIM (141-166) DMRPEIWIAQELRRIGDEFNAYYARR, hBAD (103-127) NLWAAQRYGRELRRMSDEFVDSFKK. For long-term storage at -80 °C, peptides were resuspended in water and aliquoted such that no tube was thawed more than once.

#### Intact-MS

5 μM Bcl-2 or Mcl-1 was ionized by electrospray into the Waters G2-S Synapt Quadrupole-Ion Mobility Separation-Time Of Flight Mass Spectrometer using a modified nanospray stage and key parameters noted in Table 2 at a flowrate of 6 μl/min. To prepare for ESI-MS, the protein was buffer exchanged using Slide-A-Lyzer™ MINI Dialysis Devices (2 mL, 10 kDa MWCO, Thermo Scientific™) into HPLC grade 100 mM NH_4_CH_3_COO pH 7.

**Table 2.** Synapt G2-S Parameters

| Parameter | Bcl-2 | Mcl-1 | HDX |
| --- | --- | --- | --- |
| Capillary Voltage (kV) | 3 | 3 | 3 |
| Source Temperature (°C) | 120 | 120 | 100 |
| Sampling Cone (V) | 150 | 150 | 25 |
| Cone Gas (L/Hr) | 70 | 70 | 50 |
| Nanoflow (N<sub>2</sub>, mL/min) | 2 | 2 | 2 |
| Purge Gas (N<sub>2</sub>, mL/Hr) | 600 | 600 | 600 |
| Trap Gas (Argon, mL/min) | 6 | 6 | 2 |
| Trap Cell Collision Energy (V) | 15 | 15 | 5 |
| Transfer Cell Collision Energy (V) | 10 | 10 | 5 |
| Target Enhancement (m/z) | 2250 | 2000 | None |

#### Time-resolved HDX-MS

5 μM Bcl-2/Mcl-1, or 5 μM: 30 μM Bcl-2/Mcl-1 to BH3 peptide (incubated on ice for 1 hour), underwent HDX using a kinetic mixer discussed previously^16,18^. This enabled time-resolved HDX of 1, 2, 4, and 18 seconds corresponding to inner-capillary pullback of 2, 5, 10, and 50 mm at protein and 100% D_2_O flowrates of 2 μL/min. 10% CH_3_COOH was injected at 16 μL/min to maintain a constant HDX quenching pH of 2.4 after the reaction and during proteolysis. Pepsin (porcine gastric mucosa, Sigma-Aldrich) was cross-linked in-house onto NHS-activated agarose (Pierce™, Thermo Fisher). The proteolytic chamber was constructed in-house using poly(methyl methacrylate) (PMMA) etched with a CO2 laser (VersaLaser) and affixed with a 0.2 μm pore-size frit upstream of the ESI emitter. Pepsin generated peptides were identified using ProteinLynx Global Server (Waters) after LC-MS/MS analysis with the Orbitrap Elite Hybrid Ion Trap-Orbitrap Mass Spectrometer. The deuterium uptake of peptides were analyzed using the G2-S Synapt (Waters) IMS-MS and processed by MS Studio.^26^

A maximal sequence coverage of 92% and 94% was obtained for Mcl-1 and Bcl-2, respectively. However, due to spectral overlap from digestion products of BH3 peptides, some sequence coverage was lost, resulting in 75% for Bcl2+Bim, 90% for Bcl2+Bid, 79% for Bcl2+Bad, 62% for Bcl2+Noxa, 84% for Mcl1+Bim, 81% Mcl1+Bad, 85% for Mcl1+Bid, and 92% for Mcl1+Noxa. Where possible, Expasy FindPept was used to identify peptides (by MS1) to make up for loss of redundancy (e.g., 137-150 in Bcl-2+Bad).^27^

For a given data set to be accepted, Bradykinin 2-9 (PPGFSPFR, Sigma Aldrich), was spiked in as an HDX timepoint control peptide. If Bradykinin uptake differences between the bound and unbound samples fell within the error (a statistically insignificant difference), this meant that there was no significant variation between how the two states were prepared and analyzed, and thus any changes that did occur were accurate.

#### Homology Modeling with SWISS-MODEL

Homology models were constructed using the AutoModel function in SWISS-MODEL^23^ of the following sequences: Bcl-2 Chimera MAHAGRTGYDNREIVMKYIHYKLSQRGYEW-DAGDDVEENRTEAPEGTESEPVVHLTLRQAGDDFSRRYRRDFAEMSSQLHLTPFTARGRFATVVEELFRDGVNWGRIVAFFEFGGVMCVESVNREMSPLVDNIALWMTEYLNRHLHTWIQDNGGWDAFVELYGPSMRHHHHHH; Mcl-1 GSGSDELYRQSLEIISRYLREQAT-GAKDTKPMGRSGATSRKALETLRRVGDGVQRNHETAFQGMLRKLDIKNEDDVKSLSRVMIHVFSDGVTNWGRIVTLISFGAFVAKHLKTINQESCIEPLAESITDVLVRTKRDWLVKQRGWDGFVEFFHVEDLEGG; tBid DSESQEDIIRNIARHLAQVGDSMDRSIPPGLV; Bim(EL) AEPADMRPEIWIAQELRRIGDEFNAYYARRVFL; Bad APPNLWAAQRYGR-ELRRMSDEFVDSFKKGLP; Noxa ARAPAELEVECATQLRRFGDKLNFRQKLLNLI. Structures were analyzed using PyMol to identify protein-peptide intermolecular interactions. Please see Table 3 for details.

**Table 3.** SWISS-MODEL Results and Quality

| Homology<br>Model | Template | Sequence<br>Identity (%) | Global<br>Model<br>Quality | QMEAN<br>Where 0.0 is<br>native-like. | QMEANDi<br>sCo Global |
| --- | --- | --- | --- | --- | --- |
| Bcl-2 and Bid | PDB 4QVE | 72.68 | 0.70 | -0.67 | 0.70 ± 0.06 |
| Bcl-2 and Bim | PDB 4QVF | 71.91 | 0.68 | -1.58 | 0.71 ± 0.07 |
| Bcl-2 and Bad | PDB 2BZW | 64.17 | 0.73 | -1.50 | 0.72 ± 0.06 |
| Mcl-1 and<br>Noxa | PDB 2NLA | 91.26 | 0.78 | -1.65 | 0.79 ± 0.07 |

## Supplementary material description

Sequence information; HDX peptide coverage list; Ion Mobility Data for 10+ charge state; Raw HDX uptake data.

## Abbreviations and Symbols

Bcl-2: B-cell lymphoma 2 protein
BH: Bcl-2 Homology domain
ESI-MS: Electrospray Ionization Mass
HDX-MS: Hydrogen-Deuterium Exchange Mass Spectrometry
IMS: Ion Mobility Spectrometry
Mcl-1: Induced myeloid leukemia cell differentiation protein
MOMP: Mitochondrial Outer Membrane Permeabilization
PDB: protein data bank
RMSD: root mean squared deviation
tBid: truncated Bid

## ACKNOWLEDGEMENT

This work was supported by the Natural Sciences and Engineering Research Council (NSERC) of Canada Collaborative Research and Development (CRD) (CRDPJ 485321-15) Program, the Discovery Program (RGPIN 480432) and the National Institute of General Medical Sciences (R35 GM119840) of the National Institutes of Health (BCD).

## AUTHOR CONTRIBUTIONS

**Esther Wolf:** Conceptualization (supporting); investigation (lead); writing – original draft (lead); Formal Analysis (lead). **Cristina Lento:** Writing-review and editing (supporting); Formal analysis (supporting); supervision (supporting). **Jinyue Pu:** Formal Analysis (supporting). **Bryan Dickinson**: Conceptualization (supporting); Writing – original draft (supporting); writing – review and editing (supporting); funding acquisition (supporting); supervision (supporting). **Derek Wilson**: Conceptualization (lead); writing – original draft (supporting); Writing - review and editing (lead); funding acquisition (lead); supervision (lead)

## SUPPLEMENTARY

1. Primary Sequences for Bcl2 & Mcl1
2. % Coverage and Redundancy
3. Peptide Lists
4. IMS of 10+ Apo vs. Bid Complex
5. Kinetic Plot Data for each state (8 figures)

