## Supplemental Tables and Figures for "Innate Conformational Dynamics Drive Binding Specificity in Anti-Apoptotic Proteins Mcl-1 and Bcl-2"

**Supplementary Material**

**Table S1.** Sequences of Bcl-2 and Mcl-1

|  | amino acid # | |  | amino acid # | | |
| --- | --- | --- | --- | --- | --- | --- |
| Bcl2 Chimera | | WT Bcl-2 (Bcl-XL in red) | Mcl1 (Untagged) | | WT Mcl-1 | GST-Mcl1 |
|  | 173 total |  |  | 160aa |  |  |
| M | 1 | 1 | *G* | 1 |  | 225 |
| A | 2 | 2 | *S* | 2 |  | 226 |
| H | 3 | 3 | *G* | 3 |  | 227 |
| A | 4 | 4 | *S* | 4 |  | 228 |
| G | 5 | 5 | D | 5 | 172 | 229 |
| R | 6 | 6 | E | 6 | 173 | 230 |
| T | 7 | 7 | L | 7 | 174 | 231 |
| G | 8 | 8 | Y | 8 | 175 | 232 |
| Y | 9 | 9 | R | 9 | 176 | 233 |
| D | 10 | 10 | Q | 10 | 177 | 234 |
| N | 11 | 11 | S | 11 | 178 | 235 |
| R | 12 | 12 | L | 12 | 179 | 236 |
| E | 13 | 13 | E | 13 | 180 | 237 |
| I | 14 | 14 | I | 14 | 181 | 238 |
| V | 15 | 15 | I | 15 | 182 | 239 |
| M | 16 | 16 | S | 16 | 183 | 240 |
| K | 17 | 17 | R | 17 | 184 | 241 |
| Y | 18 | 18 | Y | 18 | 185 | 242 |
| I | 19 | 19 | L | 19 | 186 | 243 |
| H | 20 | 20 | R | 20 | 187 | 244 |
| Y | 21 | 21 | E | 21 | 188 | 245 |
| K | 22 | 22 | Q | 22 | 189 | 246 |
| L | 23 | 23 | A | 23 | 190 | 247 |
| S | 24 | 24 | T | 24 | 191 | 248 |
| Q | 25 | 25 | G | 25 | 192 | 249 |
| R | 26 | 26 | A | 26 | 193 | 250 |
| G | 27 | 27 | K | 27 | 194 | 251 |
| Y | 28 | 28 | D | 28 | 195 | 252 |
| E | 29 | 29 | T | 29 | 196 | 253 |
| W | 30 | 30 | K | 30 | 197 | 254 |
| D | 31 | 31 | P | 31 | 198 | 255 |
| A | 32 | 32 | M | 32 | 199 | 256 |
| G | 33 | 33 | G | 33 | 200 | 257 |
| D | 34 | 34 | R | 34 | 201 | 258 |
| D | 35 | 35 | S | 35 | 202 | 259 |
| V | 36 | 36 | G | 36 | 203 | 260 |
| E | 37 | 37 | A | 37 | 204 | 261 |
| E | 38 | 38 | T | 38 | 205 | 262 |
| N | 39 | 39 | S | 39 | 206 | 263 |
| R | 40 | 40 | R | 40 | 207 | 264 |
| T | 41 | 41 | K | 41 | 208 | 265 |
| E | 42 | 42 | A | 42 | 209 | 266 |
| A | 43 | 43 | L | 43 | 210 | 267 |
| P | 44 | 44 | E | 44 | 211 | 268 |
| E | 45 | 45 | T | 45 | 212 | 269 |
| G | 46 | 46 | L | 46 | 213 | 270 |
| T | 47 | 47 | R | 47 | 214 | 271 |
| E | 48 | 48 | R | 48 | 215 | 272 |
| S | 49 | 49 | V | 49 | 216 | 273 |
| E | 50 | 50 | G | 50 | 217 | 274 |
| P | 51 | 91 | D | 51 | 218 | 275 |
| V | 52 | 92 | G | 52 | 219 | 276 |
| V | 53 | 93 | V | 53 | 220 | 277 |
| H | 54 | 94 | Q | 54 | 221 | 278 |
| L | 55 | 95 | R | 55 | 222 | 279 |
| T | 56 | 96 | N | 56 | 223 | 280 |
| L | 57 | 97 | H | 57 | 224 | 281 |
| R | 58 | 98 | E | 58 | 225 | 282 |
| Q | 59 | 99 | T | 59 | 226 | 283 |
| A | 60 | 100 | A | 60 | 227 | 284 |
| G | 61 | 101 | F | 61 | 228 | 285 |
| D | 62 | 102 | Q | 62 | 229 | 286 |
| D | 63 | 103 | G | 63 | 230 | 287 |
| F | 64 | 104 | M | 64 | 231 | 288 |
| S | 65 | 105 | L | 65 | 232 | 289 |
| R | 66 | 106 | R | 66 | 233 | 290 |
| R | 67 | 107 | K | 67 | 234 | 291 |
| Y | 68 | 108 | L | 68 | 235 | 292 |
| R | 69 | 109 | D | 69 | 236 | 293 |
| R | 70 | 110 | I | 70 | 237 | 294 |
| D | 71 | 111 | K | 71 | 238 | 295 |
| F | 72 | 112 | N | 72 | 239 | 296 |
| A | 73 | 113 | E | 73 | 240 | 297 |
| E | 74 | 114 | D | 74 | 241 | 298 |
| M | 75 | 115 | D | 75 | 242 | 299 |
| S | 76 | 116 | V | 76 | 243 | 300 |
| S | 77 | 117 | K | 77 | 244 | 301 |
| Q | 78 | 118 | S | 78 | 245 | 302 |
| L | 79 | 119 | L | 79 | 246 | 303 |
| H | 80 | 120 | S | 80 | 247 | 304 |
| L | 81 | 121 | R | 81 | 248 | 305 |
| T | 82 | 122 | V | 82 | 249 | 306 |
| P | 83 | 123 | M | 83 | 250 | 307 |
| F | 84 | 124 | I | 84 | 251 | 308 |
| T | 85 | 125 | H | 85 | 252 | 309 |
| A | 86 | 126 | V | 86 | 253 | 310 |
| R | 87 | 127 | F | 87 | 254 | 311 |
| G | 88 | 128 | S | 88 | 255 | 312 |
| R | 89 | 129 | D | 89 | 256 | 313 |
| F | 90 | 130 | G | 90 | 257 | 314 |
| A | 91 | 131 | V | 91 | 258 | 315 |
| T | 92 | 132 | T | 92 | 259 | 316 |
| V | 93 | 133 | N | 93 | 260 | 317 |
| V | 94 | 134 | W | 94 | 261 | 318 |
| E | 95 | 135 | G | 95 | 262 | 319 |
| E | 96 | 136 | R | 96 | 263 | 320 |
| L | 97 | 137 | I | 97 | 264 | 321 |
| F | 98 | 138 | V | 98 | 265 | 322 |
| R | 99 | 139 | T | 99 | 266 | 323 |
| D | 100 | 140 | L | 100 | 267 | 324 |
| G | 101 | 141 | I | 101 | 268 | 325 |
| V | 102 | 142 | S | 102 | 269 | 326 |
| N | 103 | 143 | F | 103 | 270 | 327 |
| W | 104 | 144 | G | 104 | 271 | 328 |
| G | 105 | 145 | A | 105 | 272 | 329 |
| R | 106 | 146 | F | 106 | 273 | 330 |
| I | 107 | 147 | V | 107 | 274 | 331 |
| V | 108 | 148 | A | 108 | 275 | 332 |
| A | 109 | 149 | K | 109 | 276 | 333 |
| F | 110 | 150 | H | 110 | 277 | 334 |
| F | 111 | 151 | L | 111 | 278 | 335 |
| E | 112 | 152 | K | 112 | 279 | 336 |
| F | 113 | 153 | T | 113 | 280 | 337 |
| G | 114 | 154 | I | 114 | 281 | 338 |
| G | 115 | 155 | N | 115 | 282 | 339 |
| V | 116 | 156 | Q | 116 | 283 | 340 |
| M | 117 | 157 | E | 117 | 284 | 341 |
| C | 118 | 158 | S | 118 | 285 | 342 |
| V | 119 | 159 | C | 119 | 286 | 343 |
| E | 120 | 160 | I | 120 | 287 | 344 |
| S | 121 | 161 | E | 121 | 288 | 345 |
| V | 122 | 162 | P | 122 | 289 | 346 |
| N | 123 | 163 | L | 123 | 290 | 347 |
| R | 124 | 164 | A | 124 | 291 | 348 |
| E | 125 | 165 | E | 125 | 292 | 349 |
| M | 126 | 166 | S | 126 | 293 | 350 |
| S | 127 | 167 | I | 127 | 294 | 351 |
| P | 128 | 168 | T | 128 | 295 | 352 |
| L | 129 | 169 | D | 129 | 296 | 353 |
| V | 130 | 170 | V | 130 | 297 | 354 |
| D | 131 | 171 | L | 131 | 298 | 355 |
| N | 132 | 172 | V | 132 | 299 | 356 |
| I | 133 | 173 | R | 133 | 300 | 357 |
| A | 134 | 174 | T | 134 | 301 | 358 |
| L | 135 | 175 | K | 135 | 302 | 359 |
| W | 136 | 176 | R | 136 | 303 | 360 |
| M | 137 | 177 | D | 137 | 304 | 361 |
| T | 138 | 178 | W | 138 | 305 | 362 |
| E | 139 | 179 | L | 139 | 306 | 363 |
| Y | 140 | 180 | V | 140 | 307 | 364 |
| L | 141 | 181 | K | 141 | 308 | 365 |
| N | 142 | 182 | Q | 142 | 309 | 366 |
| R | 143 | 183 | R | 143 | 310 | 367 |
| H | 144 | 184 | G | 144 | 311 | 368 |
| L | 145 | 185 | W | 145 | 312 | 369 |
| H | 146 | 186 | D | 146 | 313 | 370 |
| T | 147 | 187 | G | 147 | 314 | 371 |
| W | 148 | 188 | F | 148 | 315 | 372 |
| I | 149 | 189 | V | 149 | 316 | 373 |
| Q | 150 | 190 | E | 150 | 317 | 374 |
| D | 151 | 191 | F | 151 | 318 | 375 |
| N | 152 | 192 | F | 152 | 319 | 376 |
| G | 153 | 193 | H | 153 | 320 | 377 |
| G | 154 | 194 | V | 154 | 321 | 378 |
| W | 155 | 195 | E | 155 | 322 | 379 |
| D | 156 | 196 | D | 156 | 323 | 380 |
| A | 157 | 197 | L | 157 | 324 | 381 |
| F | 158 | 198 | E | 158 | 325 | 382 |
| V | 159 | 199 | G | 159 | 326 | 383 |
| E | 160 | 200 | G | 160 | 327 | 384 |
| L | 161 | 201 |  |  |  |  |
| Y | 162 | 202 |  |  |  |  |
| G | 163 | 203 |  |  |  |  |

**Table S2.** % Sequence Coverage and Redundancy

| State | % Coverage | Redundancy |
| --- | --- | --- |
| Bcl2 Only | 75 | 2.28 |
| Bcl2/Bid | 69 | 2.22 |
| Bcl2/Bim | 61 | 1.67 |
| Bcl2/Bad | 62 | 1.76 |
| Bcl2/Noxa | 55 | 1.93 |
| Mcl1 Only | 92 | 2.11 |
| Mcl1/Bid | 85 | 1.54 |
| Mcl1/Bim | 84 | 2.05 |
| Mcl1/Bad | 77 | 1.71 |
| Mcl1/Noxa | 92 | 2.11 |

**Table S3.** Mcl1 Peptide List

| Mcl-1 |  |  |  |  |  |  |
| --- | --- | --- | --- | --- | --- | --- |
| m/z | z | m | Drift Time | Sequence | Start | Stop |
| 676.8288 | 2 | 1351.642 | 3.255 | SDELYRQSLEI | ^172 | 181 |
| 589.8311 | 2 | 1178.655 | 2.604 | EIISRYLRE | 180 | 188 |
| 409.2286 | 2 | 816.4415 | 1.682 | TGAKDTKP | 191 | 198 |
| 904.4791 | 1 | 903.4713 | 7.161 | KPMGRSGAT | 197 | 205 |
| 821.4367 | 3 | 2461.287 | 3.038 | QATGAKDTKPMGRSGATSRKALET | 189 | 212 |
| 644.5959 | 4 | 2574.352 | 2.713 | QATGAKDTKPMGRSGATSRKALETL | 189 | 213 |
| 581.2994 | 3 | 1741.884 | 2.007 | RRVGDGVQRNHETAF | 214 | 228 |
| 686.676 | 3 | 2057.004 | 2.333 | RRVGDGVQRNHETAFQGM | 214 | 231 |
| 447.2531 | 4 | 1785.992 | 2.224 | LRKLDIKNEDDVKSL | 232 | 246 |
| 710.071 | 3 | 2128.201 | 3.201 | LRKLDIKNEDDVKSLSRV | 232 | 249 |
| 672.377 | 3 | 2015.12 | 2.93 | RKLDIKNEDDVKSLSRV | 233 | 249 |
| 703.3295 | 2 | 1405.652 | 3.309 | MIHVFSDGVTNW | 250 | 261 |
| 637.8085 | 2 | 1274.609 | 2.875 | IHVFSDGVTNW | 251 | 261 |
| 682.3511 | 3 | 2045.036 | 2.604 | MIHVFSDGVTNWGRIVTL | 250 | 267 |
| 638.6748 | 3 | 1914.01 | 2.333 | IHVFSDGVTNWGRIVTL | 251 | 267 |
| 708.3834 | 3 | 2122.127 | 2.55 | DGVTNWGRIVTLISFGAFVA | 256 | 275 |
| 709.9061 | 2 | 1417.797 | 3.689 | ISFGAFVAKHLKT | 268 | 280 |
| 735.89 | 2 | 1470.773 | 3.58 | VAKHLKTINQESC | 274 | 286 |
| 861.428 | 1 | 861.428 | 6.835 | SCIEPLAE | 285 | 292 |
| 667.3611 | 3 | 2000.071 | 2.441 | VRTKRDWLVKQRGWDG | 299 | 314 |
| 537.5432 | 4 | 2147.149 | 2.17 | VRTKRDWLVKQRGWDGF | 299 | 315 |
| 1002.437 | 1 | 1002.439 | 8.354 | FHVEDLEGG | 319 | 327 |

**Table S4.** Bcl2 Peptide List

| Bcl-2 |  |  |  |  |  |  |
| --- | --- | --- | --- | --- | --- | --- |
| m/z | z | m | Drift Time | Sequence | Start | Stop |
| 449.5257 | 3 | 1345.554 | 2.062 | AHAGRTGYDNRE | 2 | 13 |
| 507.7625 | 4 | 2027.019 | 2.007 | IVMKYIHYKLSQRGYE | 14 | 29 |
| 812.7342 | 3 | 2435.179 | 2.984 | VEENRTEAPEGTESEPVVHLTL | **91 | 97 |
| 907.7771 | 3 | 2720.308 | 3.309 | NRTEAPEGTESEPVVHLTLRQAGDD | *91 | 103 |
| 389.7483 | 2 | 777.4809 | 1.682 | PVVHLTL | 91 | 97 |
| 511.7341 | 2 | 1021.453 | 2.17 | TLRQAGDDF | 96 | 104 |
| 434.9001 | 3 | 1301.677 | 1.573 | FSRRYRRDF | 104 | 112 |
| 573.3018 | 3 | 1716.882 | 2.496 | SSQLHLTPFTARGRF | 116 | 130 |
| 434.9091 | 3 | 1301.704 | 1.953 | HLTPFTARGRF | 120 | 130 |
| 583.3159 | 2 | 1164.616 | 2.984 | LTPFTARGRF | 121 | 130 |
| 503.7474 | 2 | 1005.479 | 2.224 | LFRDGVNW | 137 | 144 |
| 447.2057 | 2 | 892.3957 | 1.736 | FRDGVNW | 138 | 144 |
| 695.3737 | 2 | 1388.732 | 3.201 | FRDGVNWGRIVA | 138 | 149 |
| 512.9453 | 3 | 1535.812 | 2.116 | FRDGVNWGRIVAF | 138 | 150 |
| 768.8913 | 2 | 1535.767 | 3.7 | FRDGVNWGRIVAFF | 138 | 151 |
| 789.3725 | 2 | 1576.729 | 3.58 | CVESVNREMSPLVD | 158 | 171 |
| 829.4379 | 2 | 1656.86 | 3.852 | SVNREMSPLVDNIAL | 161 | 175 |
| 534.2533 | 3 | 1599.736 | 2.007 | WMTEYLNRHLHT | 176 | 187 |
| 800.887 | 2 | 1599.758 | 4.12 | MTEYLNRHLHTW | 177 | 188 |
| 654.7814 | 2 | 1307.547 | 2.93 | WIQDNGGWDAF | 188 | 198 |
| 469.2087 | 4 | 1872.803 | 2.007 | VELYGPSMRHHHHHH | 199 | 207-6His |

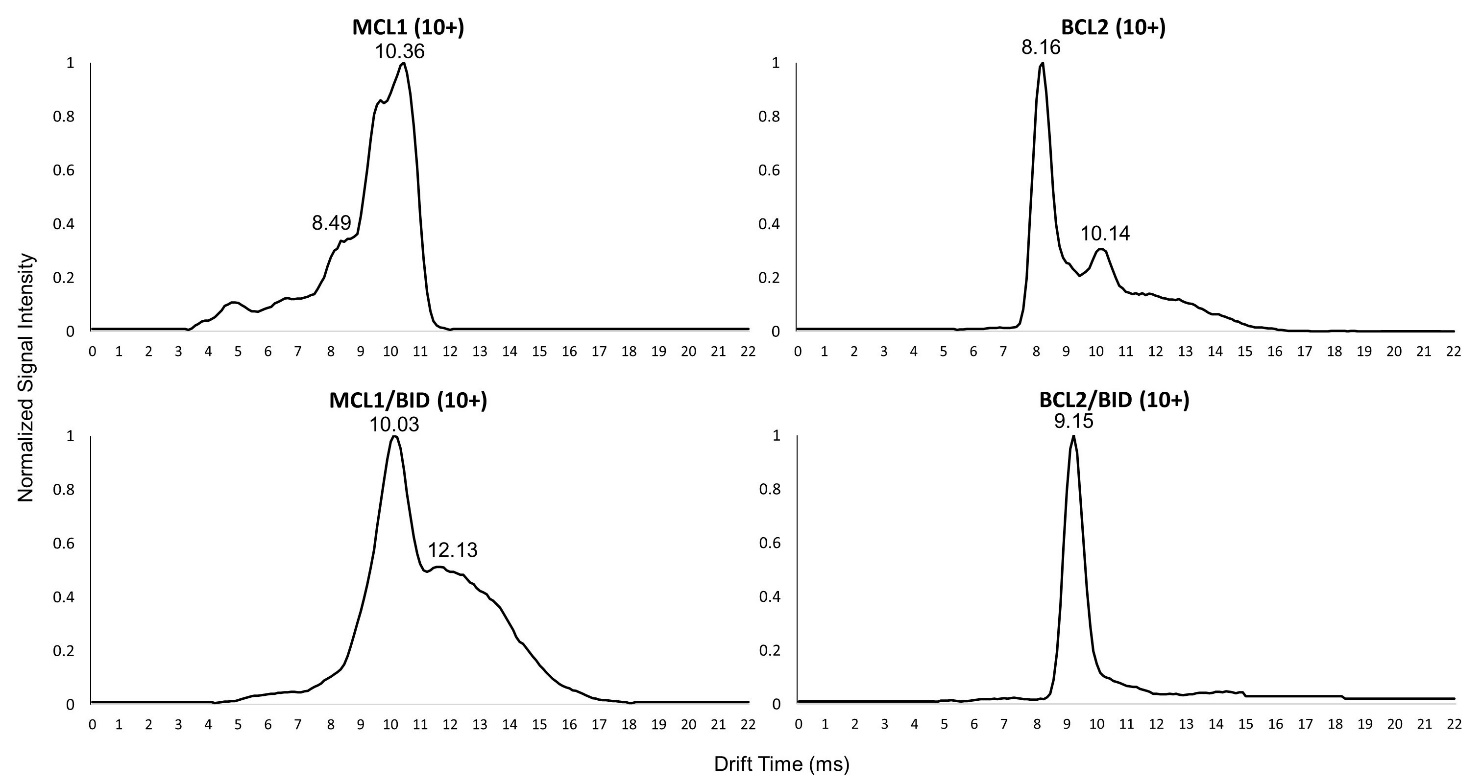

**Figure S1.** Ion Mobility Chromatograms of Bcl-2 and Mcl-1. The relative signal intensity as a function of drift time (milliseconds) for Mcl-1 (top left), Bcl-2 (top right), Mcl-1 + Bid (bottom left), and Bcl-2 + Bid (bottom right) is shown for their 10+ charge state. These 10+ charge states are less abundant and lower intensity (ion count).

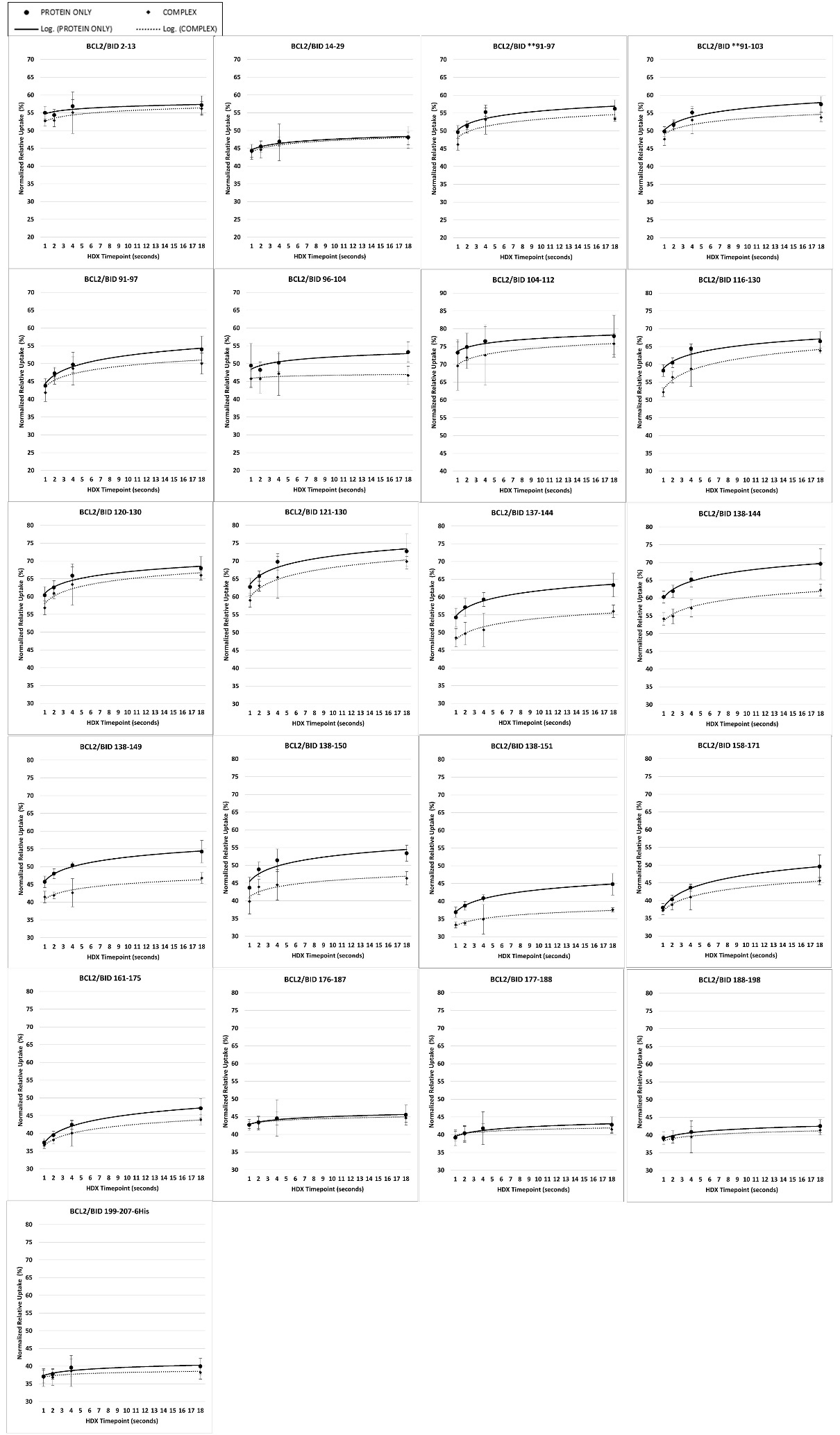

**Figure S2.** Bcl2 vs. Bcl2/Bid Deuterium Uptake Kinetics Plots.

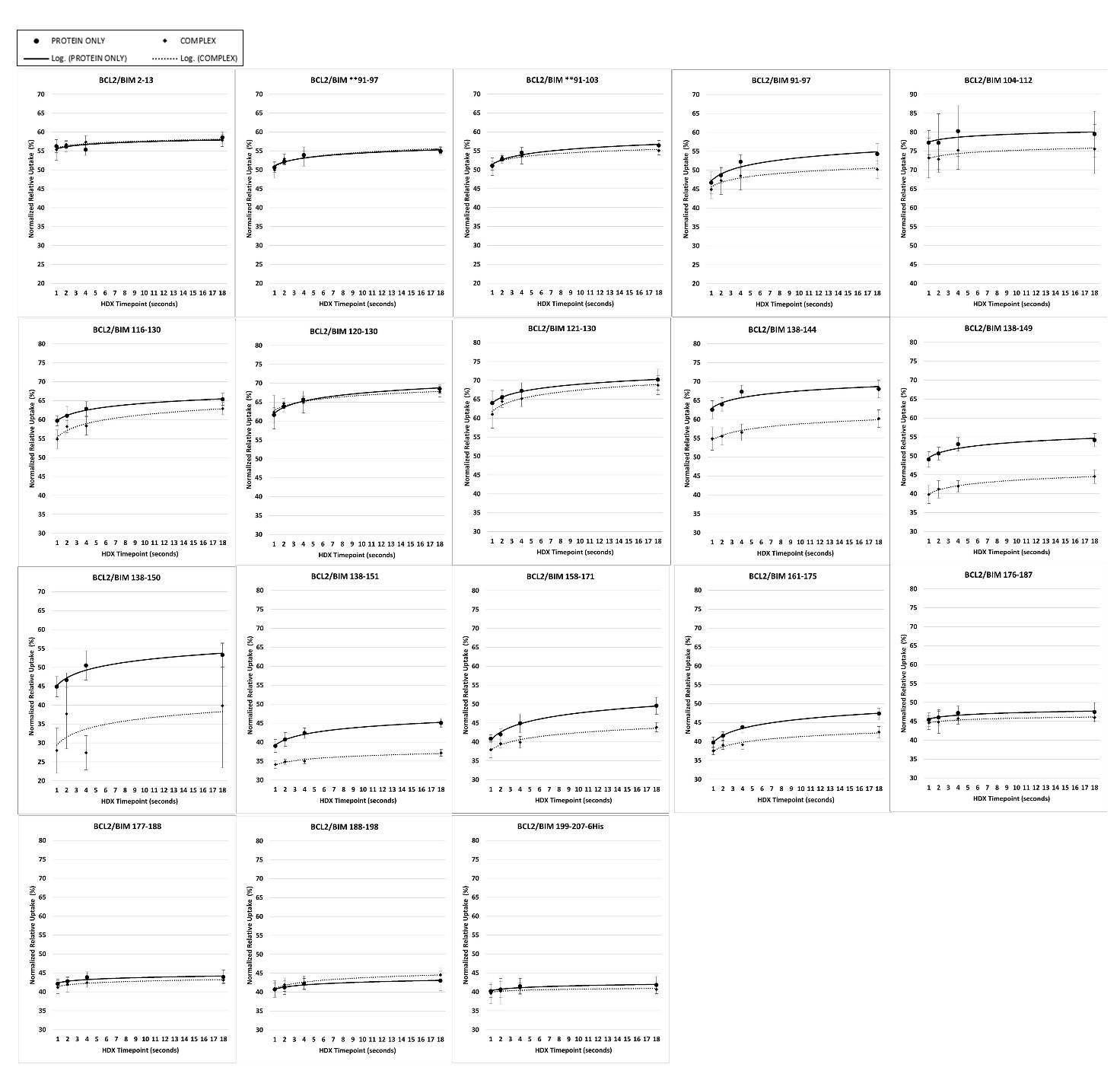

**Figure S3.** Bcl2 vs. Bcl2/Bim Deuterium Uptake Kinetics Plots.

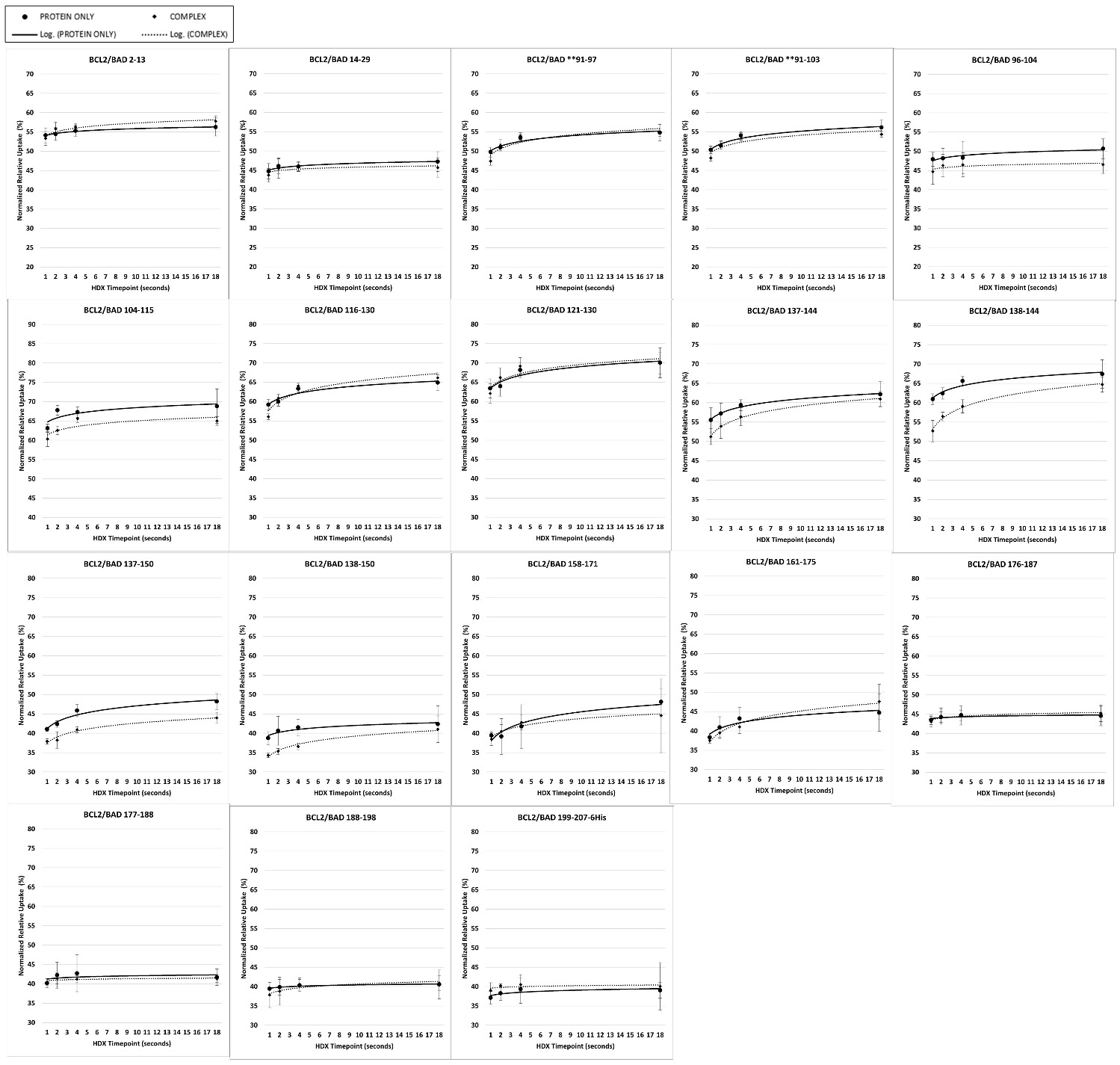

**Figure S4.** Bcl2 vs. Bcl2/Bad Deuterium Uptake Kinetics Plots.

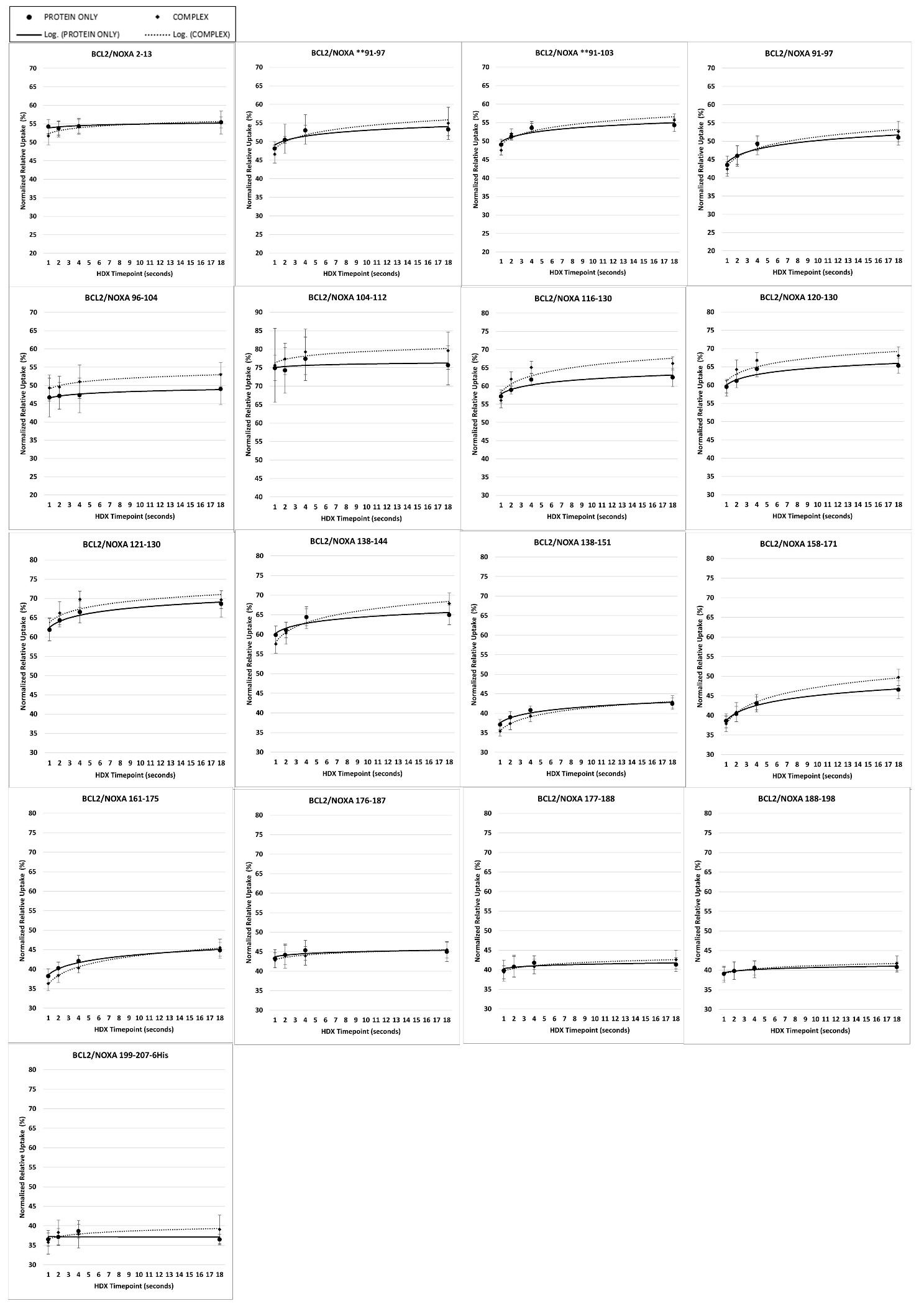

**Figure S5.** Bcl2 vs. Bcl2/Noxa Deuterium Uptake Kinetics Plots.

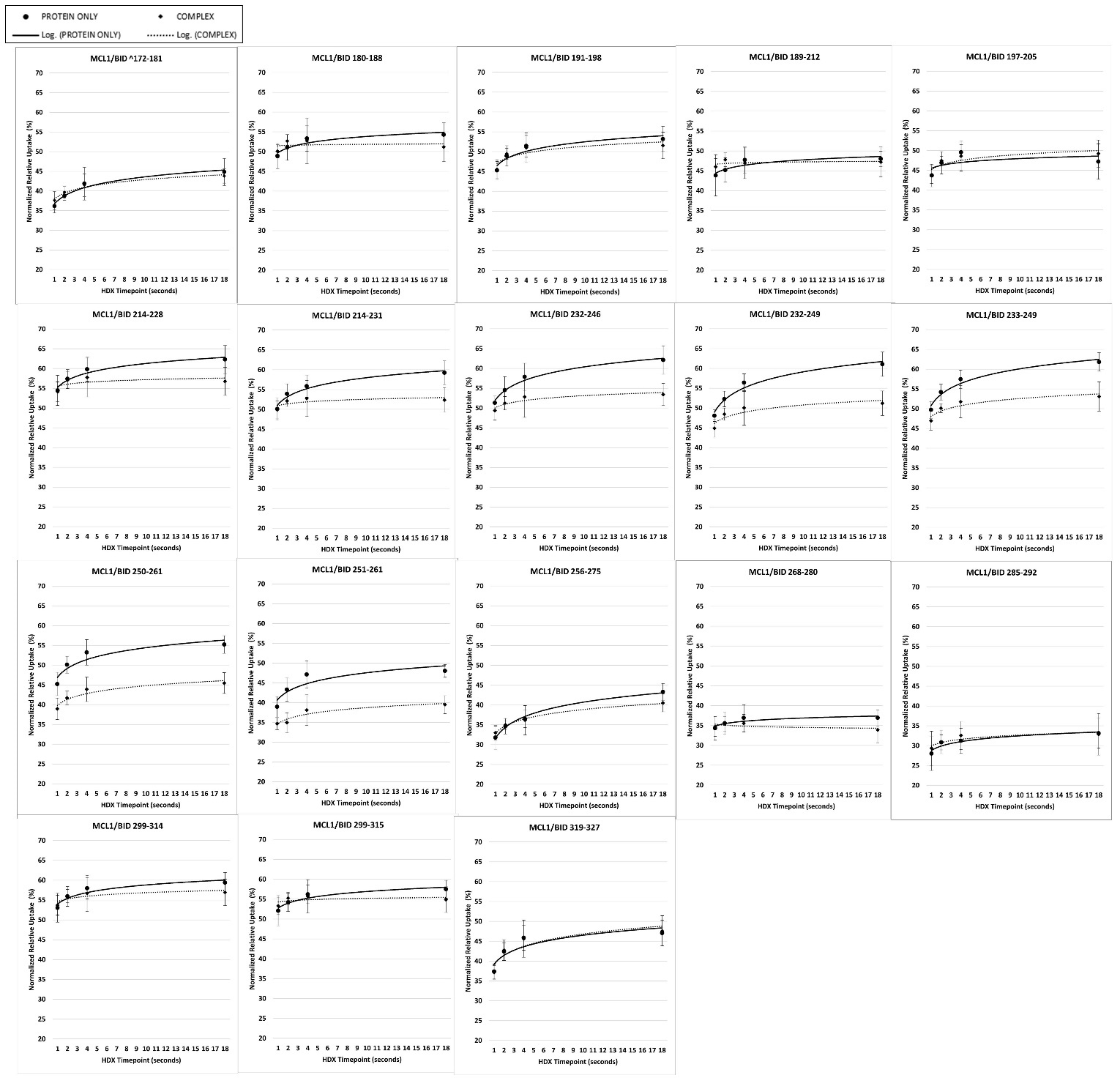

**Figure S6.** Mcl1 vs. Mcl1/Bid Deuterium Uptake Kinetics Plots.

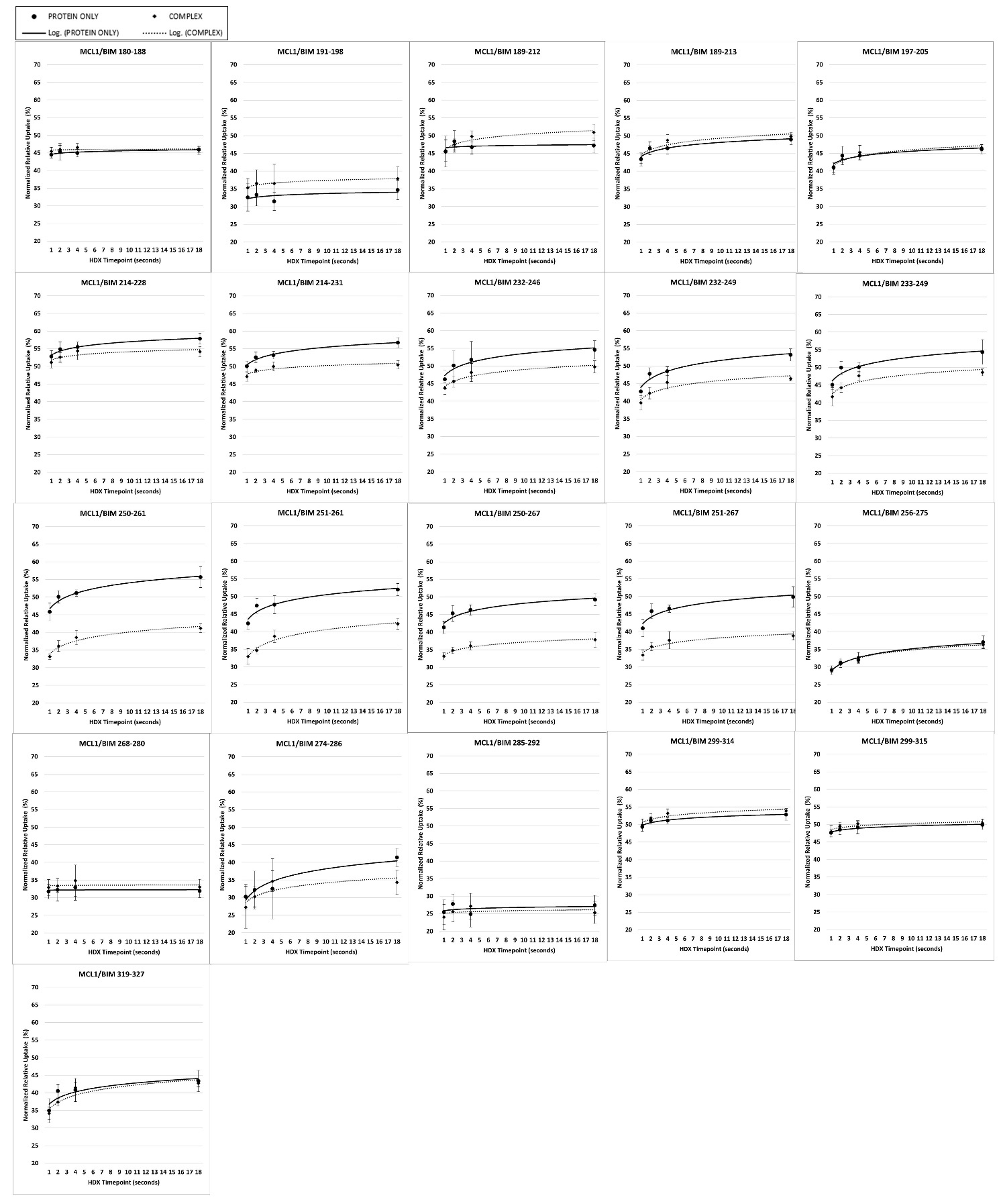

**Figure S7.** Mcl1 vs. Mcl1/Bid Deuterium Uptake Kinetics Plots.

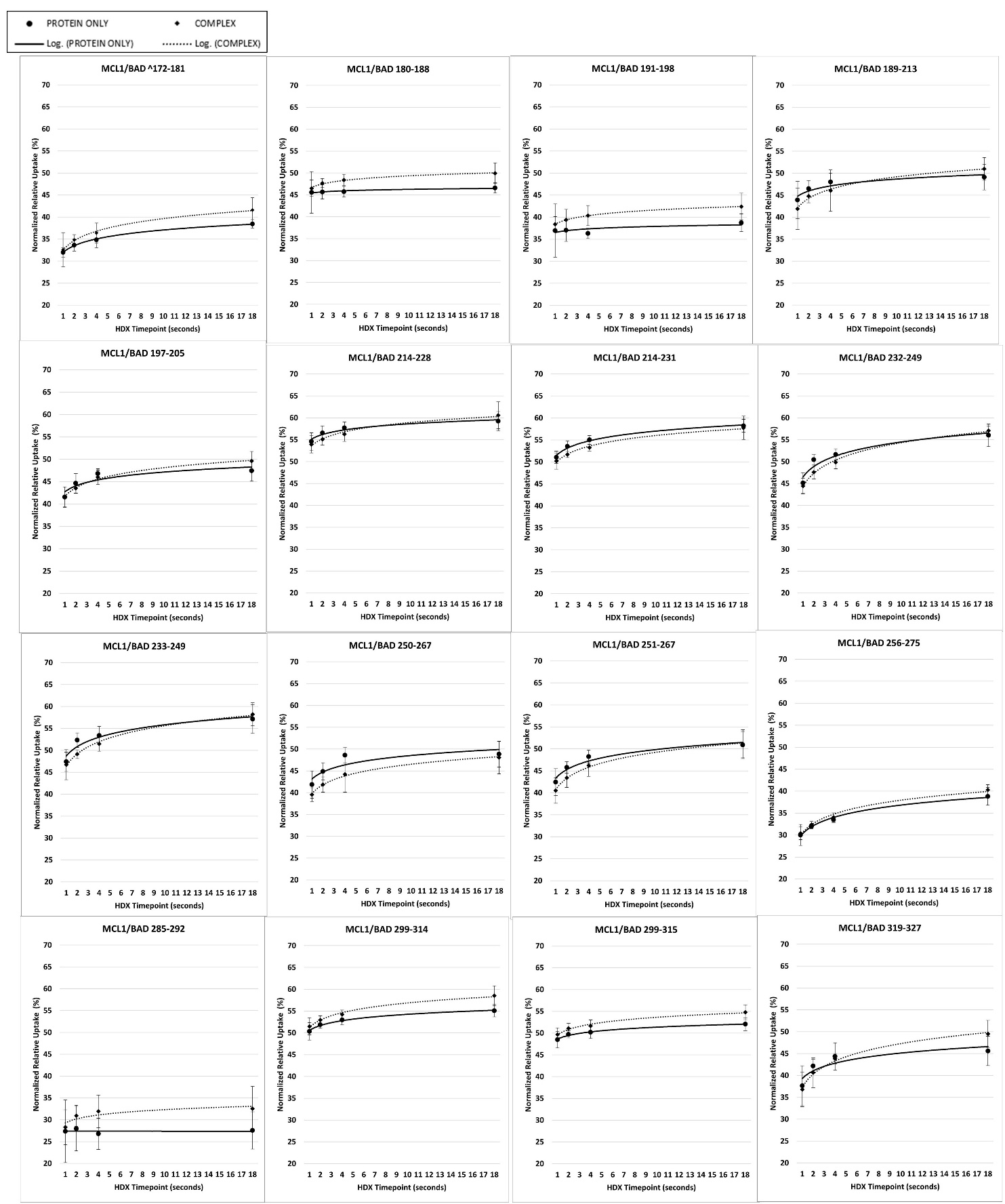

**Figure S8.** Mcl1 vs. Mcl1/Bad Deuterium Uptake Kinetics Plots.

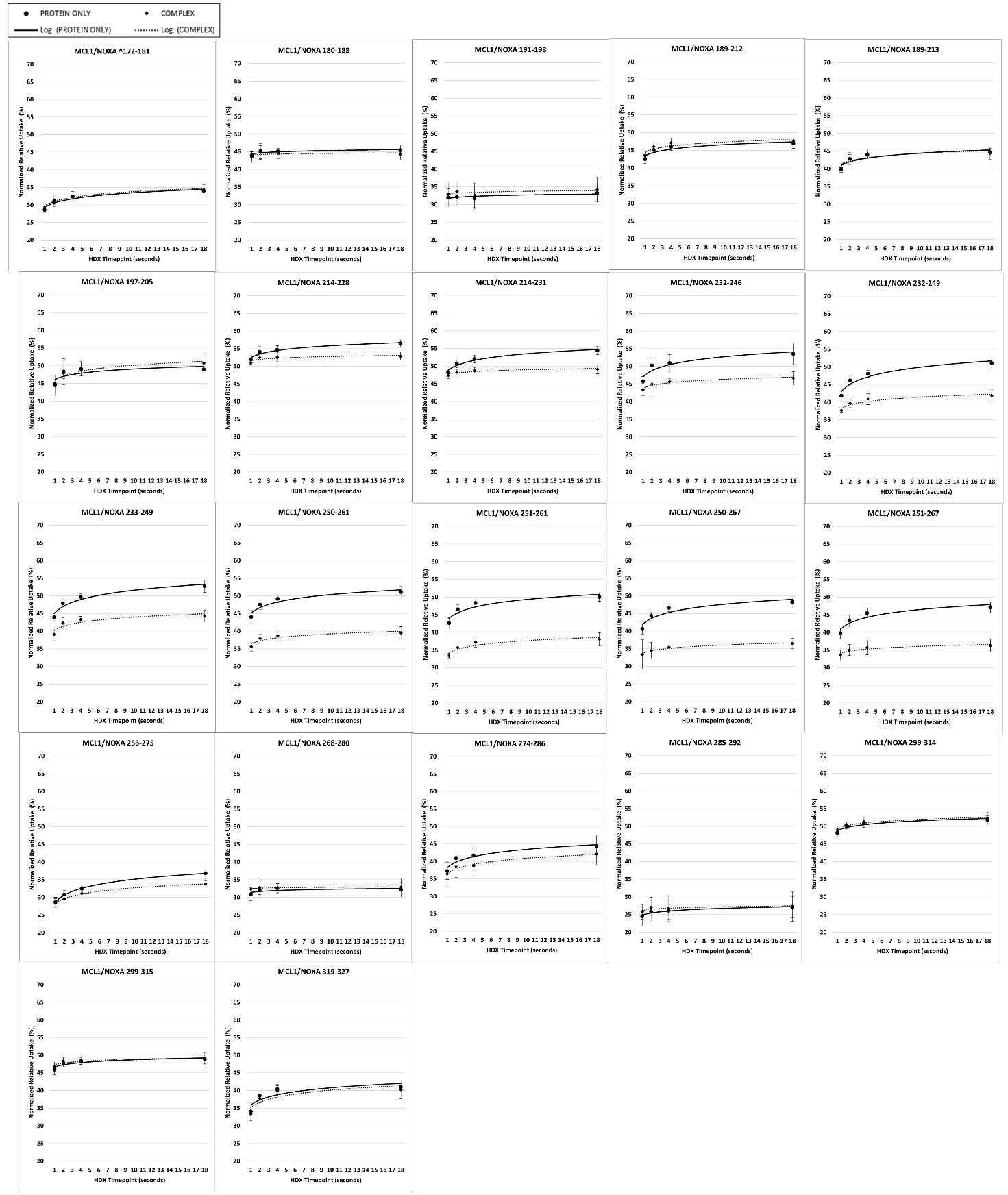

**Figure S9.** Mcl1 vs. Mcl1/Noxa Deuterium Uptake Kinetics Plots.
